# The transcriptional landscape of plant infection by the rice blast fungus *Magnaporthe oryzae* reveals distinct families of temporally co-regulated and structurally conserved effectors

**DOI:** 10.1101/2022.07.18.500532

**Authors:** Xia Yan, Bozeng Tang, Lauren S. Ryder, Dan MacLean, Vincent M. Were, Alice Bisola Eseola, Neftaly Cruz-Mireles, Andrew J. Foster, Miriam Osés-Ruiz, Nicholas J. Talbot

## Abstract

The rice blast fungus *Magnaporthe oryzae* causes a devastating disease which threatens global rice production. In spite of intense study, the biology of plant tissue invasion during blast disease remains poorly understood. Here we report a high resolution, transcriptional profiling study of the entire plant-associated development of the blast fungus. Our analysis revealed major temporal changes in fungal gene expression during plant infection. Pathogen gene expression could be classified into 10 modules of temporally co-expressed genes, providing evidence of induction of pronounced shifts in primary and secondary metabolism, cell signalling and transcriptional regulation. A set of 863 genes encoding secreted proteins are differentially expressed at specific stages of infection, and 546 were predicted to be effectors and named *MEP* (*<u>M</u>agnaporthe* <u>e</u>ffector <u>p</u>rotein) genes. Computational prediction of structurally-related *MEPs*, including the MAX effector family, revealed their temporal co-regulation in the same co-expression modules. We functionally characterised 32 *MEP* genes and demonstrate that Mep effectors are predominantly targeted to the cytoplasm of rice cells via the biotrophic interfacial complex (BIC), and use a common unconventional secretory pathway. Taken together, our study reveals major changes in gene expression associated with blast disease and identifies a diverse repertoire of effectors critical to successful infection.

## INTRODUCTION

Rice blast disease is one of the most significant constraints on rice production worldwide. Each year, despite deployment of rice varieties carrying numerous resistance genes and the extensive use of fungicides, between 10 and 30% of the rice harvest is lost to rice blast disease (Yamaguchi, 2004; Wang and Valent, 2017; Stam and McDonald, 2018; Poloni et al., 2020). Given that rice provides 23% of the calories to humankind and is a staple food for half of the world’s population, controlling rice blast in a sustainable way, would constitute a major contribution to global food security. Rice blast is caused by the filamentous fungus *Magnaporthe oryzae* (synonym of *Pyricularia oryzae*) (Zhang et al., 2016) which has evolved the ability to breach the tough outer cuticle of rice leaves and invade living plant tissue. *M. oryzae* can also infect a wide range of grass hosts causing diseases such as wheat blast, for instance, an emerging threat to wheat production (Urashima. et al., 1993; Islam et al., 2016; Inoue et al., 2017; Latorre et al., 2022).

The blast fungus undergoes a series of morphogenetic transitions during plant infection, involving cellular differentiation to develop infection cells called appressoria on the leaf surface, followed by penetration and rapid proliferation of invasive hyphae within rice cells, facilitated by active suppression of plant immunity. Like many of the most important crop pathogens, rice blast fungus is a hemibiotroph, which means that it actively grows in living plant tissue during its initial biotrophic phase of development, but later causes plant cell death when it develops necrotrophically, utilising nutrients released from dead plant cells to enable the fungus to sporulate from necrotic disease lesions.

So far, more than 1600 *M. oryzae* genes have been functionally characterised by targeted gene deletion or mutagenesis– greater than 10% of the rice blast genome (Foster et al., 2021). These studies have provided insight into the biology of blast disease, especially the ability of the fungus to form appressoria on the leaf surface, but there are many aspects of the biology of blast disease that are not well understood, even after such intense study. We do not know, for example, how the blast fungus is able to invade living plant tissue so rapidly, overcoming host defences, colonising new plant cells, and spreading long distances throughout rice leaves. To be such an efficient invader of plant cells, *M. oryzae* must be able to adapt successfully to the host environment– sequestering carbon and nitrogen sources to fuel its growth –while evading recognition by the host, but how this is achieved is not clear. The blast fungus also induces profound changes in the organisation of plant cells, including extensive membrane biogenesis, changes in cytoskeletal configuration and perturbation of cell-to-cell communication. How these processes are orchestrated and regulated by the invading fungus also remains largely unknown. One of the principal reasons for our current lack of understanding is that there have been very few investigations which have taken a holistic view of plant infection– in contrast to gene functional studies –attempting to understand the progression of blast disease and the major temporal changes in pathogen physiology.

In this study, we set out to define the transcriptional landscape of rice blast infection. Our aim was to identify major changes in pathogen gene expression from the moment of initial inoculation of plants until development of disease symptoms and, in particular, to use this information to identify the full repertoire of fungal effector proteins deployed by the fungus.

Effectors are secreted proteins that target components of the plant immune system to suppress host defense and enable proliferation of the pathogen (Jones and Dangl, 2006; Kamoun, 2006; Lo Presti et al., 2015). In addition to suppression of plant immunity, effectors may target cell signalling and metabolic processes to help facilitate fungal invasive growth (Zhai et al., 2022). In *M. oryzae,* effectors target extracellular processes such as chitin-triggered immunity that operate in the apoplast (Mentlak et al., 2012), or intracellular processes such as perturbation of reactive oxygen species generation (Liu and Zhang, 2021), targeted protein degradation (Park et al., 2012) or re-programming host transcription (Kim et al., 2020). Intracellular effectors accumulate in a membrane-rich plant structure called the biotrophic interfacial complex (BIC), which appears necessary for their delivery into plant cells (Kankanala et al., 2007), and four cytoplasmic effectors have been shown to be secreted by an unusual Golgi-independent mechanism (Giraldo et al., 2013). A sub-set of *M. oryzae* effectors are recognised by rice immune receptors leading to disease resistance. The interaction of such avirulence effectors (Avrs) with cognate NLR receptors has also helped reveal their likely intracellular targets, especially in cases where specific effector-binding domains have become integrated into NLRs (Cesari et al., 2013; Maqbool et al., 2015; De la Concepcion et al., 2018). Several sequence-un-related Avr effectors possess a conserved protein fold and are termed MAX effectors (de Guillen et al., 2015). Structural modelling of the predicted secreted proteome of *M. oryzae* has recently identified large sets of structurally-related proteins (Seong and Krasileva, 2021a), although their role in blast disease is not yet known.

We reasoned that comprehensive transcriptional profiling would provide a means by which the landscape of blast disease could be systematically analysed at a holistic level and the true effector repertoire of *M. oryzae* revealed. Transcriptomic studies have, for instance, provided key insights into physiologically complex states, such as tumorigenesis (Wingrove et al., 2019), embryonic development (He et al., 2020) and host immunity (Bjornson et al., 2021), as well as fungal-plant interactions (Copley et al., 2017; Zeng et al., 2017; Lanver et al., 2018). We recognised that in the *M. oryzae*-rice interaction there have been previous attempts to define global patterns of gene expression, using methods such as microarrays and super-SAGE analysis or, more recently, by RNA-seq (Mosquera et al., 2009; Chen et al., 2012; Kawahara et al., 2012; Soanes et al., 2012; Dong et al., 2015; Sharpee et al., 2017; Shimizu et al., 2019). These studies have, however, suffered from the lower resolution of previous methodologies, failing to detect gene expression from the low fungal biomass during early stages of plant infection. Relatively poor coverage of fungal gene expression changes have therefore been reported, with only the most abundantly expressed fungal genes identified. More recent studies using RNA-seq analysis have benefited from the dynamic range and resolution of deep sequencing, but these investigations have focused, almost exclusively, on a single time point during infection, leading to assumptions regarding the progress of disease that have severe limitations. Moreover, most transcriptomic studies of *M. oryzae* have generated only superficial coverage of fungal gene expression changes, because of the type of inoculation methods carried out and the relatively poor rates of blast infection observed.

In this project, we set out to overcome the major limitations of previous studies by carrying out a comprehensive time-course of rice blast disease using different infection protocols and rice hosts to enhance the efficiency of blast infection. We reasoned that by infecting rice cultivars of varying blast susceptibility, coupled with using different inoculation methods, we could optimise the number of fungal genes analysed and thereby generate deeper insights into the transcriptional landscape of plant infection. We performed RNA-seq analysis of *M. oryzae* strain Guy11 infecting two rice cultivars of different susceptibility, either in spray infections or using leaf drop inoculation of attached rice leaves. We report the global pattern of fungal gene expression at 8 time-points from 0h (the time of spore inoculation) until full blast symptom expression at 144h. In this way we were able to define 10 modules of temporally co-expressed fungal genes and define physiological, metabolic, and gene regulatory networks represented by each module. We identified 863 secreted protein-encoding genes that are differentially-regulated during plant infection, many of which are predicted to encode effectors. The effector repertoire of *M. oryzae* and its temporal expression dynamics are therefore much greater in complexity than previously recognised. Strikingly, we also found that effector candidates predicted to be structurally conserved (Seong and Krasileva, 2021) are temporally co-expressed during the biotrophic growth. Using live cell imaging and gene functional analysis, we report the cell biological features of these effectors and their specific positioning during infection. When considered together, our findings provide new insight into the biology of blast disease and the complexity of the deployed effector repertoire.

## RESULTS

To determine the transcriptional landscape of rice blast disease, we first selected two rice cultivars differing in their susceptibility to blast. CO39 is a dwarf *indica* variety of rice, moderately susceptible to blast, that has been used as a host in many gene functional studies of *M. oryzae* (Talbot et al., 1993a; Chauhan et al., 2002; Zhu et al., 2004). Moukoto, meanwhile, is a *japonica* rice variety, highly susceptible to blast disease and resulting in large, coalescing disease lesions (Yoshida et al., 2009). We inoculated 21-day-old rice seedlings by spray infection with 1 x 10^5^ conidia mL^-1^, as well as carrying out leaf drop infections in which a 20 μl drop of a suspension of 1 x 10^6^ conidia mL^-1^ was placed on the surface of a rice leaf which remained attached to a 21-day-old rice seedling. The compatible *Magnaporthe oryzae* strain Guy11 was used for all infections and rapid invasion of new rice cells could be observed in using live cell imaging of a GFP-expressing strain of Guy11, between 40-48h after inoculation, as shown in Figure 1A and Supplemental Movie 1. All infection experiments were repeated three times with *M. oryzae* cultures and rice seedlings of the same age and always inoculated at the same time of day to control for circadian effects (Figure 1B). The inoculated leaf area was collected for sample extraction at 8 different times which are key stages for disease development; 0h (uninfected), 8h, 16h, 24h, 48h, 72h, 96h and 144h post-infection as depicted in Figure 2. These time points cover all morphogenetic transitions associated with appressorium development, appressorium-mediated penetration, biotrophic growth, transpressorium-dependent cell-to-cell movement, fungal proliferation in rice tissue, the switch to necrotrophic growth and fungal sporulation (Cruz-Mireles et al., 2021). Using Illumina sequencing of mRNA libraries extracted from rice tissue, we generated 4.37 billion reads from all samples (Supplemental Dataset 1). All data is freely accessible through ENA Accession Number PRJEB45007. Kraken 2 was used to identify *M. oryzae* and rice reads from the mixed transcriptome (Wood et al., 2019). We classified reads either as rice or the rice blast fungus and focus here on reads mapping to the annotated *M. oryzae* reference genome (Dean et al., 2005). By utilizing the highly susceptible Moukoto, we were able to show a greater number of *M. oryzae* sequence reads at earlier stage of infection and throughout disease progression compared with CO39 infections, as shown in Figure 1C. We observed, however, that leaf drop infections were significantly enriched in fungal reads in the mixed transcriptome analysis. The greater number of fungal reads probably reflects a higher proportion of rice cells infected by the fungus in leaf drop samples compared to spray inoculations, so the transcriptome data are likely to be more representative of the infected state. By contrast, in spray inoculations many rice cells remain uninfected throughout and will only respond at a distance from the site of infection (Figure 1C and Supplemental Figure 1A). During infection, we observed a 3-fold increase in the proportion of *M. oryzae* reads between 24 hpi and 48 hpi in leaf drop infections (Supplemental Dataset 1), consistent with the steep increase in fungal biomass that accompanies invasion of neighbouring cells during this 24h period (Figure 1C and Figure 2B). Principal Component Analysis (PCA) using Transcripts Per Kilobase Million (TPM) highlighted the reproducibility between biological replicates of each sample (Supplemental Figure 1B) and also provided evidence of a significant change in gene expression between early stages (8 hpi-24 hpi) of infection and later stages (48 hpi-144 hpi), which occurs irrespective of inoculation method (Supplemental Figure 2A).

**Figure 1.**
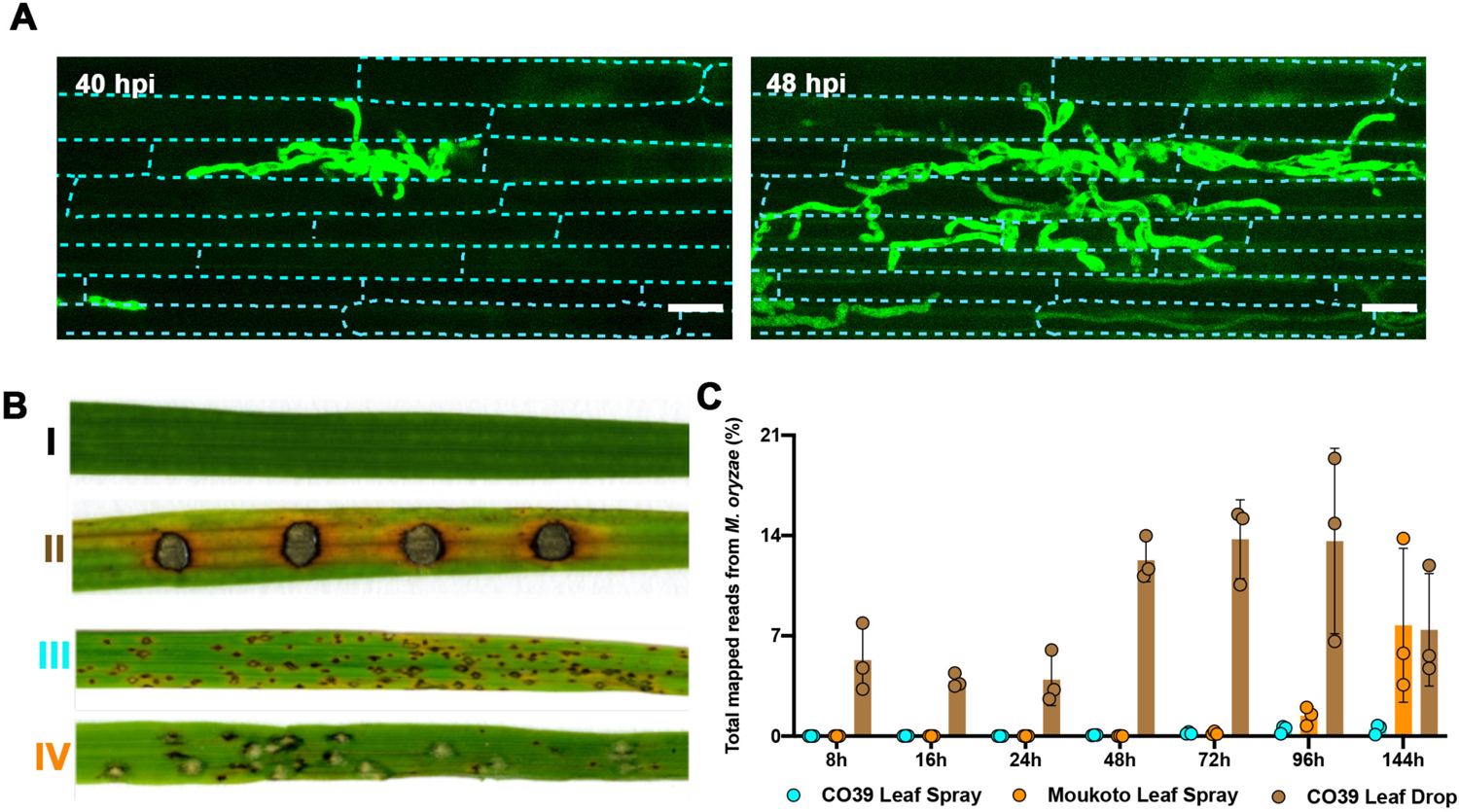
Transcriptional profile analysis of a time course of plant infection by the rice blast fungus *M. oryzae.* Rice infections were carried out using two distinct inoculation methods and two cultivars differing in relative susceptibility to blast. **(A)** Micrographs of rice tissue of cultivar CO-39 inoculated with *M. oryzae* Guy11 expressing free GFP driven by the constitutive promoter RP27 at 40 h post inoculation (hpi) and 48 hpi to show progression of tissue invasion. Individual plant cells are outlined by the cyan dotted line. Scale bars = 20 µm. **(B)** Comparison of rice blast disease symptoms 6 days post-inoculation using either leaf drop infection or spray infection on rice cultivars with varying host susceptibility. I: moderately susceptible rice cultivar CO-39 inoculated with water control; II: leaf drop infection of rice CO-39 with Guy11; III: spray infection of rice CO-39 with Guy11; IV: spray infection of highly susceptible rice cultivar Moukoto with Guy11. **(C)** Graph depicting proportion of fungal transcripts in the plant and pathogen mixed transcriptome (CO-39 Leaf Spray, Moukoto Leaf Spray and CO-39 Leaf Drop correspond to inoculation methods).

**Figure 2.**
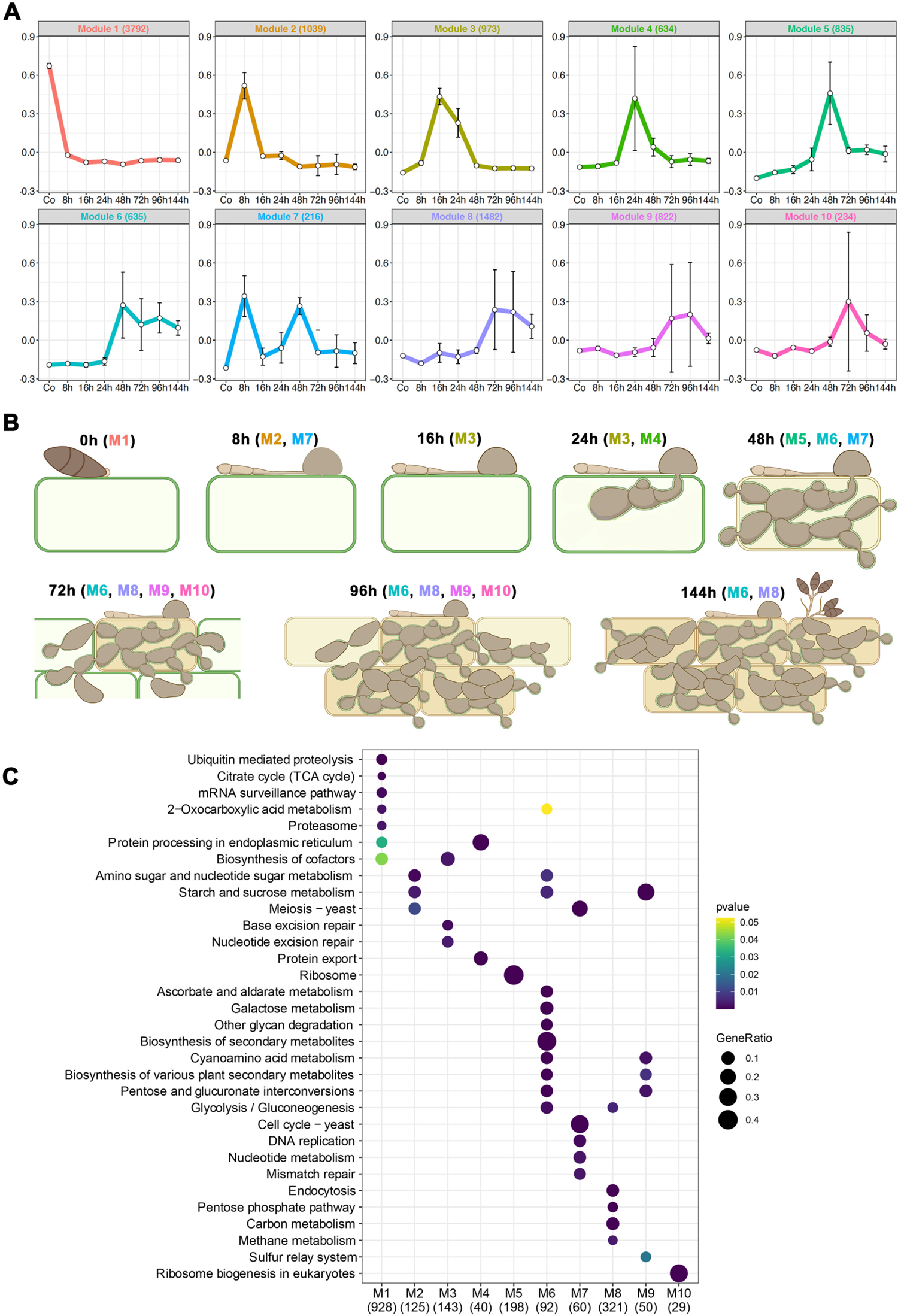
Temporal co-expression analysis reveals 10 modules of pathogen gene expression during rice blast infection. Analysis of co-expressed pathogen genes during rice blast disease development. **(A)** Weighted correlation network analysis (WGCNA) identified 10 co-expressed modules during a time-course of infection-related-development and plant infection (Modules 1-10). The representative eigengene is shown for each module **(B)** Schematic representation of each stage of rice blast disease development and corresponding WGCNA co-expression modules. Blast-infected rice tissue samples were collected at 0 hpi, 8 hpi, 16 hpi, 24 hpi, 48 hpi, 72 hpi, 96 hpi and 144 hpi. **(C)** KEGG enrichment analysis of genes in each WCGNA module using clusterProfiler reveals over-represented physiological functions during blast disease development.

### Temporal dynamics of gene expression during rice blast disease

To define the temporal sequence of fungal gene expression changes during rice infection, we analysed gene expression across the time course using weighted gene co-expression network analysis (WGCNA) (Zhang and Horvath, 2005; Langfelder and Horvath, 2008). This analysis identified 10 modules of co-expressed genes, for which the expression pattern of a representative eigengene is shown in Figure 2. Modules ranged in size from 216 to 3792 temporally co-expressed genes (Figure 2A and Supplemental Dataset 2). Genes in color-coded modules are highly expressed at the developmental stages shown in Figure 2B. Module (M) 1, for example, represents 3792 genes that show peak expression in conidia and are then down-regulated following appressorium-mediated penetration. These include genes involved in fungal growth, conidiation and spore germination, such as *LEU1* (Tang et al., 2019; Que et al., 2020), *SOM1*(Yan et al., 2011) and *YPT7* (Liu et al., 2015), all previously reported to be necessary for sporulation, as well as genes implicated in appressorium morphogenesis, such as the osmosensor-encoding gene *SHO1* (Liu et al., 2011), the appressorium turgor sensor kinase gene *SLN1* (Zhang et al., 2010; Ryder et al., 2019), and the *RGS* family of genes that regulate G-protein signalling (Zhang et al., 2011). The septins are also predominantly classified in M1 (*SEP3*, *SEP4* and *SEP6*) and M2 (*SEP5*) consistent with their vital role in appressorium re-polarisation (Dagdas et al., 2012).

Genes associated with appressorium-mediated infection were predominantly represented by M2, which shows peak expression at 8hpi (Figure 2B), and contains 1039 genes highly expressed during the pre-penetration phase of fungal growth. These include *PTH11* (Sweigard et al., 1998), *BUF1* (Valent et al., 1991), *MAGB* (Liu and Dean, 1997) and *ZNF1* (Cao et al., 2016; Yue et al., 2016), all necessary for appressorium development and function. Initial biotrophic colonisation of rice tissue is represented by genes in M3, M4 and M5, which show peaks of gene expression at 16hpi, 24hpi and 48hpi, respectively. M3 contains *MSB2* (Liu et al., 2011), *HOX7* (Kim et al., 2009; Oses-Ruiz et al., 2021) and *MET13* (Yan et al., 2013) implicated in appressorium function, and *SEC6,* which encodes a component of the octameric exocyst complex, that assembles in a septin-dependent manner during plant infection (Gupta et al., 2015). M4 consists of 634 genes including gene functions expected during early biotrophic growth, such as *RBF1* which encodes a protein required for BIC formation (Nishimura et al., 2016), as well as many effectors (see below). M5 contains 835 genes associated with invasive growth, such as *NOXD*, the NADPH oxidase sub-unit-encoding gene (Galhano et al., 2017) as well as the *MPS1* MAPK (Xu et al., 1998), *SSB1*, *SSZ1* and *ZUO1* which are all involved in the cell wall integrity pathway (Yang et al., 2018). This is consistent with a substantial change in cell wall organisation occurring during invasive growth. M6 contains genes up-regulated in association with the increase in fungal biomass that occurs during invasive growth. The module is also enriched in genes encoding transcription factors and signalling proteins (Supplemental Data Set 2), which are predominantly uncharacterised, and may reflect the dramatic reprogramming in fungal physiology that occurs at the switch from biotrophic growth to necrotrophic growth. M7 shows genes which peak in expression during appressorium development at 8 h and then again during transpressorium development at 48h. This observation is consistent with the developmental conservation of these two infection structures, as recently highlighted (Cruz-Mireles et al., 2021). M7 contains *MST12*, for example, which encodes a transcription factor required for appressorium-mediated penetration (Park et al., 2002), *CPKA* encoding the catalytic sub-unit of cAMP-dependent protein kinase A, another key regulator of appressorium development (Mitchell and Dean, 1995), as well as the cell cycle regulator *NIM1* (Saunders et al., 2010) and peroxin-encoding gene *PEX1* (Deng et al., 2016). Their bi-modal expression profile in M7 is consistent with these functions being implicated in transpressorium morphogenesis. Finally, the switch to necrotrophic growth by *M. oryzae* is represented by M8, M9 and M10, which peak in expression after 72hpi (Figure 2B). These modules include gene functions associated with conidiogenesis, such as *SMO1*, encoding a Ras GTPase-Activating Protein, the fatty acid synthase beta subunit dehydratase *FAS1* (Sangappillai and Nadarajah, 2020) and the necrosis- and ethylene-inducing protein 1-encoding gene *NLP1* (Park et al., 2006; Fang et al., 2017; Kershaw et al., 2019).

### Invasive growth of *M. oryzae* involves specific physiological transitions

To identify physiological processes within each WGCNA module of co-expressed genes, we next carried out metabolic pathway enrichment analysis, as shown in Figure 2C. As the temporal progression of rice blast infection proceeds, several waves of gene expression could be identified. Over-representation of genes associated with regulated proteolysis, for example, are a feature of M1, consistent with the role of autophagy in appressorium maturation. There is also a switch from the tricarboxylic acid cycle (M1), to sucrose metabolism (M2, M6, M9) and the pentose phosphate pathway during invasive growth (M8). This is consistent with the Nut1/Pas1 /Asd4-regulated NADPH-dependent metabolic regulation previously reported in *M. oryzae* (Wilson et al., 2010) and the metabolic transitions that occur during fungal invasive growth (Fernandez et al., 2014). The physiological signature of biotrophy also includes evidence of rapid fungal proliferation, exhibited by cell division-associated functions in M2 and M3, as well as protein processing and export in M4, and ribosomal biogenesis in M5. There is also a very pronounced switch that is clear in M6, associated with the appearance of disease symptoms, which occurs at 72h, and the onset of necrotrophy. M6 shows over-representation of secondary metabolism-associated gene expression and plant cell wall degrading enzymes, for instance. The parallels between appressorium morphogenesis and transpressorium function are also evident by the physiological functions over-represented in M7, including cell cycle control– a key regulator of appressorium development(Saunders et al., 2010)–and associated DNA replication functions. Further secondary metabolic functions are represented in M8 and M9 during the onset of disease symptom development and conidiogenesis.

In a complementary analysis to WCGNA clustering, we defined expression profiles of all *M. oryzae* genes at each stage during infection. Fungal read counts were normalized and compared to expression in conidial mRNA using Sleuth (Pimentel et al., 2017), thereby identifying temporal changes in gene expression (log2 fold change >1, and P-adj<0.05) that occur during pathogenesis compared to the original inoculum. This analysis revealed that the largest change in *M. oryzae* gene expression occurs at 48hpi compared to the conidial transcriptome, highlighting the substantial physiological transition associated with invasive fungal growth, with 1920 genes up-regulated and 1012 down-regulated at this time point. Metabolic pathway enrichment analysis revealed expression of many growth-related functions, including cell division, translation, and regulated proteolysis, as well as the switch to carbohydrate metabolism that accompanies fungal invasive growth (Supplemental Figure 3; Supplemental Dataset 5). Differentially expressed genes that showed repression (or marked down-regulation) during each stage of pathogenic development provided evidence of a general repression of transport-associated functions, lipid and fatty acid metabolism occurs during invasive growth compared to conidia (Supplemental Figure 3). This highlights how pathogenic development requires not only up-regulation of specific gene functions, but also the orchestrated repression of many gene functions compared to the relative pluripotency of the germinating spore.

Targeted analysis of secondary metabolic functions revealed specific expression of polyketide synthase (PKS), fatty acid synthase (FAS), and cytochrome P450 mono-oxygenases during plant infection (Supplemental Figures 4-6). Conidial-specific expression of 8 PKS genes occurs prior to infection, followed by co-regulation of 4 PKS genes at 8h, 6 at 24 h and 7 at 48h, consistent with the production of specific metabolites during each stage of development (Supplemental Figure 4). This is mirrored by FAS gene expression occurring differentially during both biotrophic and necrotrophic stages of development (Supplemental Figure 5) and, strikingly, by temporal expression of cytochrome P450 genes at each timepoint with 21 genes expressed for example specifically at 24h (Supplemental Figure 6). Consistent with these temporal changes in gene expression, a targeted analysis of the 495 predicted transcription factor-encoding genes from the *M. oryzae* genome (Park et al., 2013) showed a pattern of co-expression, with large clusters of co-expressed TF genes in conidia, and at the 8h, 16h, 24h and 48h time points in particular (Supplemental Figure 7). A clear transition in transcriptional regulation occurs after 48h, with most transcription factor-encoding genes from the earlier biotrophic phase of growth being repressed after this time, accompanied by the expression of a completely separate group of regulators between 72h and 144 h during disease symptom development. We conclude that major regulatory changes in gene expression, associated with switches in both primary and secondary metabolism occur during plant infection and the transition from biotrophic proliferation to necrotrophic growth.

### A large family of *M. oryzae* effector genes is differentially expressed during rice infection

We next focused on defining the expression of *M. oryzae* genes predicted to encode secreted proteins, including potential effector candidates. Secreted proteins were predicted using SignalP from data sets of each inoculation method and strain-cultivar interaction. This revealed 68 candidate effector genes in the spray inoculation of CO39, 467 effector candidates in the spray inoculation of Moukoto and 847 putative effectors in the leaf drop infection of CO39 (Figure 3 and Supplemental Datasets 3-5), consistent with the increased resolution obtained by using a highly susceptible rice cultivar and leaf drop infections. We then investigated the temporal expression profile of predicted effectors and classified them into each WGCNA module (Figure 3A). This revealed distinct patterns of gene expression, particularly during the initial stages of gene expression (M1-M5), with particular over-representation of effector candidates in M4 and M5. The expression of well-characterized effector-encoding genes corresponded to each module. M3, for example, includes *NIS1* which suppresses Bak1/Bik1-dependent PAMP-triggered immunity (Mentlak et al., 2012; Irieda et al., 2019), the apoplastic effector gene *SLP1* which suppresses chitin-triggered immunity (Mentlak et al., 2012) and *MoSVP1*, encoding an effector required for appressorium-mediated penetration (Shimizu et al., 2019).

**Figure 3.**
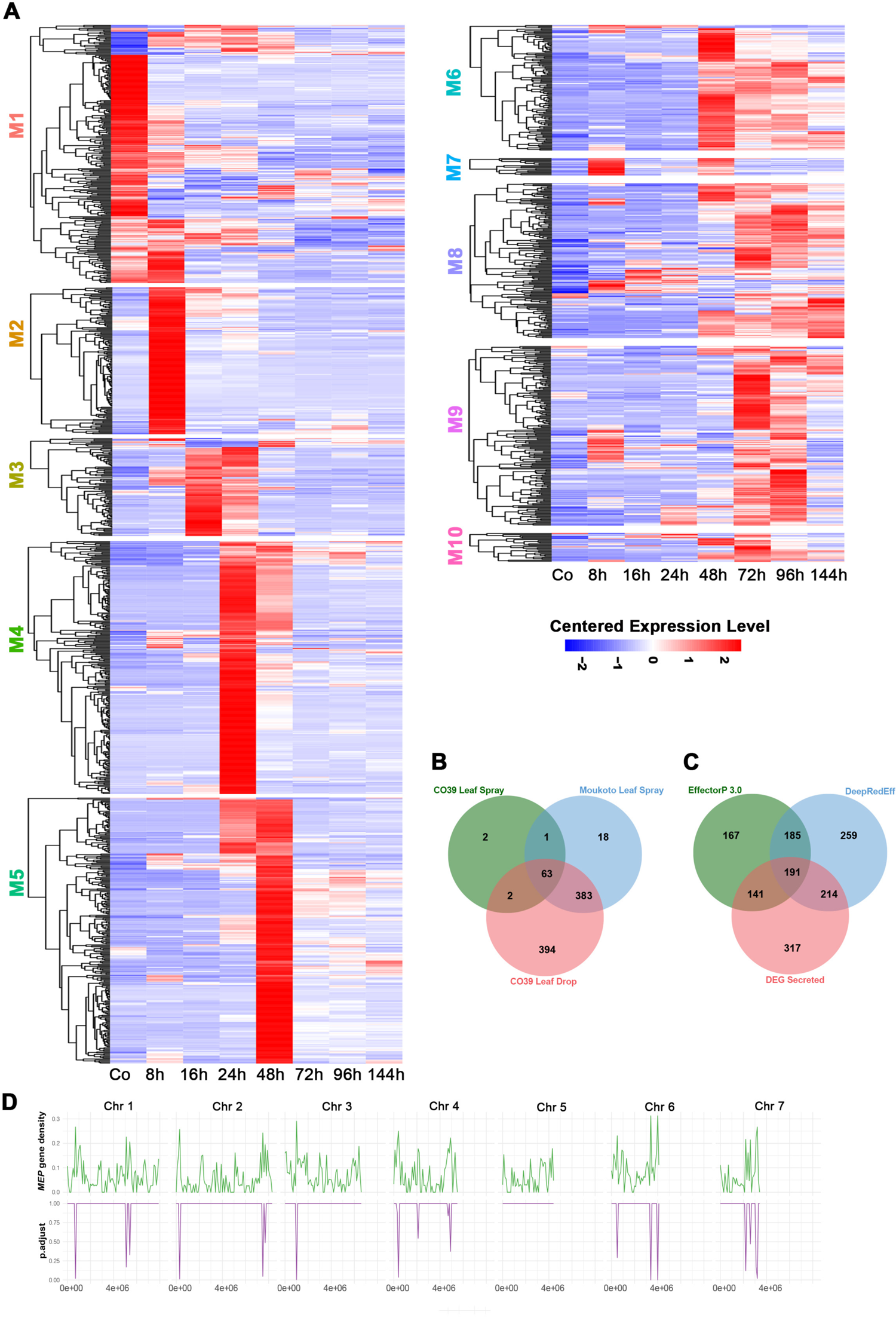
Stage-specific temporal expression of the *M. oryzae* secretome during rice infection. **(A)** Heat map showing the hierarchical clustering of pathogen genes encoding putatively secreted proteins from each WGCNA co-expression module. Distinct temporal patterns of secretome expression occur during biotrophic and necrotrophic development. Expression levels are shown relative to the mean expression TPM (transcript per kilobase million) value across all stages. **(B)** Venn diagram showing comparison of differentially expressed secreted protein-encoding genes from three different inoculation experiments (CO39 Leaf Spray, Moukoto Leaf Spray, and CO39 Leaf Drop Infections). In total 68 effector candidates were identified from the CO39 leaf spray infections, 467 effector candidates from Moukoto leaf spray inoculation, and 847 effector candidates from CO39 leaf drop infections. Although leaf drop-CO39 infections revealed expression of many more differentially expressed effector candidates, 20 effector candidates were revealed only in spray inoculation experiments. **(C)** Comparison of effector predictions of host-induced secreted proteins using EffectorP 3.0 and DeepRedEff algorithms with differentially expressed signal peptide containing genes (DEG Sec). **(D)** Enrichment analysis of *MEP* gene loci on *M. oryzae* chromosomes 1-7, encoding predicted secreted proteins. The number of *MEP* genes was compared to total gene number in 200kb windows across each chromosome. Phyper was used to compute the hypergeometric distribution and obtain P-values. (Johnson et al., 1992). The resulting graph in the bottom panel shows regions of each chromosome where there is significant over-representation of *MEP* gene loci.

M4 includes the avirulence gene *ACE1*, *HEG13* which antagonizes cell death induced by Nep1 in *Nicotiana benthaniana* (Mogga et al., 2016; Guo et al., 2019), and the *BAS1*, *BAS2* and *BAS4* effectors (Bohnert et al., 2004; Mosquera et al., 2009). M5 includes *AVR-Pita1* and *BAS3* as well as the cell death-inducing protein CDIP5 (Orbach et al., 2000; Mosquera et al., 2009), while M6 includes *SPD2* which is a suppressor of cell death (Sharpee et al., 2017). Necrotrophy associated effectors are meanwhile represented in later modules, such as the necrosis inducing protein *NLP1* in M10 (Mogga et al., 2016).

We therefore defined a total of 863 putative *<u>M</u>agnaporthe* <u>e</u>ffector <u>p</u>rotein (*MEP*) genes differentially expressed during plant infection and co-expressed with many previously reported effectors (Supplemental Dataset 7). To test the hypothesis that these genes encode fungal effectors, we used two algorithms designed to predict effector-like genes, the EffectorP algorithm and the DeepRedEff algorithm (Kristianingsih and MacLean, 2021; Sperschneider and Dodds, 2022). A total of 684 genes were predicted to be effector-encoding using EffectorP 3.0 and 849 genes using the DeepRedEff algorithm, based on analysis of the entire *M. oryzae* predicted proteome (Figure 3C). In total, 546 host-induced *MEP* genes are therefore predicted to encode effectors based on at least one of the algorithms (Figure 3C). There are, however, 317 *MEP* genes predicted not to be effectors based on either algorithm. We found that 442 predicted *MEP* effectors do not contain any PFAM domains. The products of *MEP4, MEP12* and *MEP20*, however, contain a C2H2 zinc finger domain, and *MEP31* encodes a predicted secreted homeodomain protein. *MEP12* and *MEP20* were independently identified as *MoHTR1* and *MoHTR2* which encode nuclear effectors that modulate host immunity via transcriptional reprogramming (Kim et al., 2020). *MEP161* has also been described as *MoSVP1*, an *in planta*-expressed virulence effector (Shimizu et al., 2019), while *MEP11* is an ortholog of *AVR-Pik* (Yoshida et al., 2009). We conclude that *M. oryzae* strain Guy11 has at least 546 effector genes and potentially as many as 863, which we have classified as differentially expressed *MEP*s.

### *MEP* genes are unevenly distributed in chromosomes of *M. oryzae* strain Guy11

To investigate the distribution of the differentially expressed *MEP* genes in the *M. oryzae* genome, we investigated their chromosomal distribution (Supplemental Figure X). We observed the full potential repertoire of 863 *MEP* genes at loci dispersed across all 7 chromosomes of *M. oryzae*. There appeared to be enrichment in *MEP* loci in sub-telomeric regions of all chromosomes except chromosome 5, as shown in Figure 3D. *BAS4*, *MEP4*, *MEP7*, *MEP11*, *MEP13* and *MEP24*, for example, are located near telomeres and their enrichment is statistically significant (Figure 3D). There were also *MEP* genes found adjacent to one another, for example, *MEP5* and *AVRPITA* are tightly linked in the sub-telomeric region of one arm of chromosome 6 (Supplemental Figure 8).

### Structurally conserved *M. oryzae* effectors are temporally co-regulated during plant infection

Fungal effectors seldom show sequence similarity, or homology to known proteins, but there is increasing evidence that they may share structural conservation. Indeed, structural biology has contributed significantly to our understanding of effector evolution and function (Franceschetti et al., 2017). In *M. oryzae* the MAX effectors (Magnaporthe Avrs and ToxB like), for example, are not related by sequence similarity but share a common β-sandwich fold shared by effectors from the wheat tan spot pathogen *Pyrenophora tritici-repentis* (de Guillen et al., 2015). MAX effectors include *M. oryzae* AVR1-CO39 and AVR-PikD which interact with heavy metal-associated (HMA) integrated domains in the immune receptors by which they are perceived during incompatible interactions (De la Concepcion et al., 2018; Guo et al., 2018).

Recently, a genome-wide computational structural analysis of the secreted proteome of *M. oryzae* has been reported (Seong and Krasileva, 2021a), taking advantage of the development of *de novo* folding algorithms, such as Alphafold (Jumper et al., 2021). A total of 1854 secreted proteins were classified into 905 structure clusters, 740 of which are represented by single proteins (Seong and Krasileva, 2021a). We decided to determine the relationship between the structurally conserved families reported (Seong and Krasileva, 2021b) and the predicted *MEP* effector repertoire. We found that 863 *MEP* proteins fall into 366 predicted structure clusters (Supplemental Data Set 7). This includes 38 predicted MAX effectors including *MEP3*, *MEP11* (*AVRPikD*), *MEP19* and *MEP27*, which are classified in structure cluster 8. We then selected the 10 largest structurally conserved clusters and analysed their distribution in each WGCNA module, as shown in Figure 4. In most cases the structurally conserved proteins were expressed throughout pathogenesis, but some striking patterns were observed. First of all, the MAX effector group (cluster 8), were over-represented in M4 and M5, along with structure cluster 7 proteins, which are predicted ADP-ribose transferases (ARTs), proposed to act as effectors (Seong and Krasileva, 2021a). This was a very distinct pattern from all of the other structurally conserved groups, which were much more evenly distributed. The other pattern observed was some over-representation of proteins predicted to encode hydrolases (cluster 1), glucosidases (cluster 2), and glycosyl hydrolases (clusters 4 and 6) in M8 and M9, associated with necrotrophy. We conclude that structurally conserved effector candidates are temporally co-expressed during plant infection.

**Figure 4.**
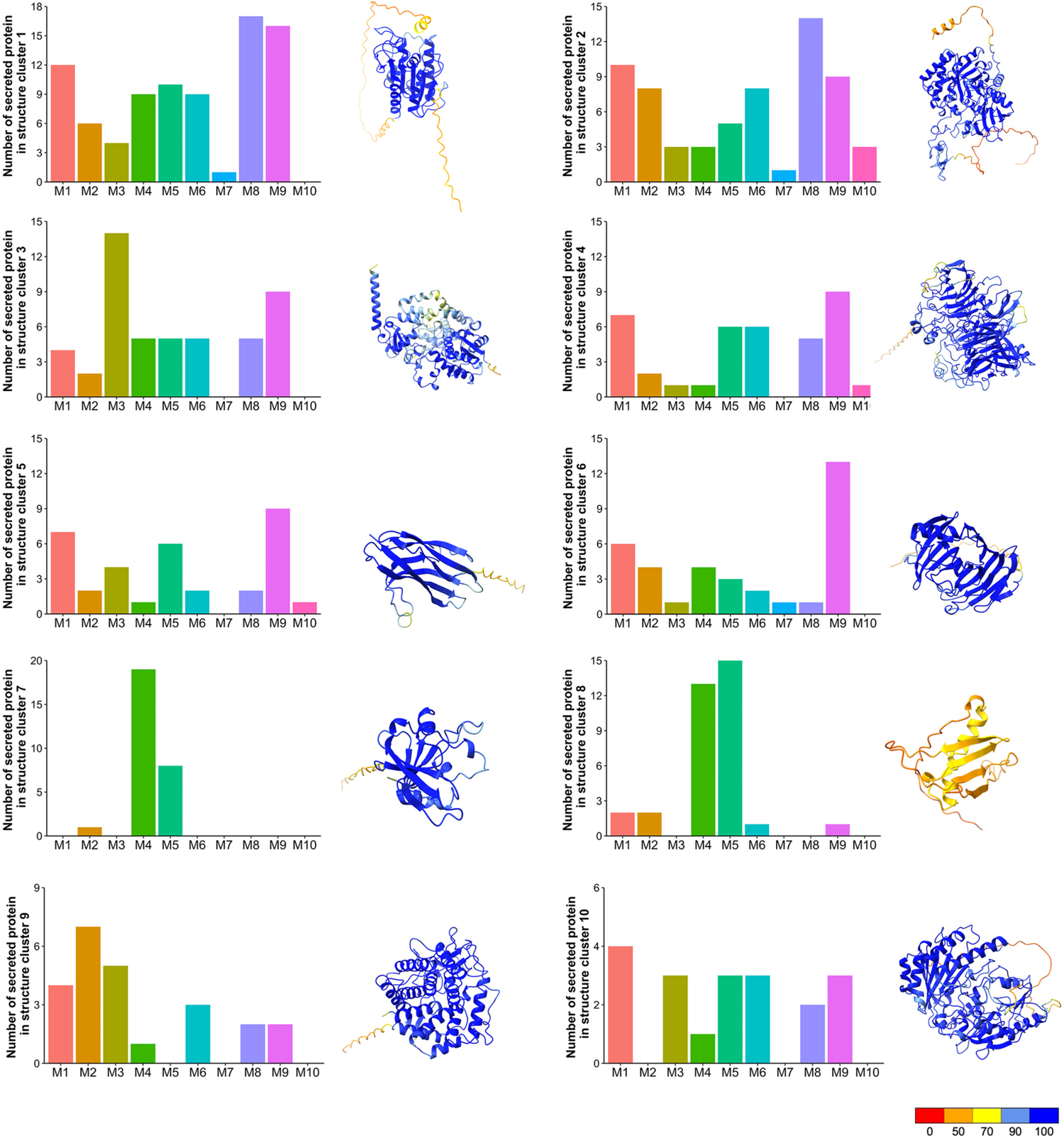
Structurally conserved *M. oryzae* effectors are temporally co-expressed during biotrophic invasive growth. AlphaFold and ChimeraX platforms were used to predict the three-dimensional structures of predicted secreted proteins of *M.oryzae.* Structure clusters were then analysed against each WGCNA module. Bar charts represent the number of predicted secreted proteins identified in each predicted structure cluster (clusters 1-10), that are classified within each WGCNA co-expression module during pathogenesis. Ribbon diagrams show a representative predicted protein structure for each *M. oryzae* structure cluster. These are MGG_09840, which contains a cutinase domain for structure cluster 1; MGG_04534 contains a chitin recognition protein domain for structure cluster 2; MGG_07497 contains a cytochrome P450 domain for structure cluster 3; MGG_05785 (INV1/BAS113) contains a glycosyl hydrolase family 32 domain for structure cluster 4; MGG_02347 (Nis1) for structure cluster 5; MGG_08537 contains a glycosyl hydrolase family 12 domain for structure cluster 6; for structure cluster; MGG_00230 is a predicted ADP-ribose transferase containing a heat-labile enterotoxin alpha chain domain for structure cluster 7; MGG_12426 is a predicted MAX effector and homologue of AvrPib for structure cluster 8; MGG_04305 contains a glycosyl hydrolase family 88 domain for structure cluster 9; MGG_00276 contains berberine and berberine-like domains for structure cluster 10. Protein structures were predicted by AlphaFold. and ChimeraX was used to visualise the protein structure using PDB files generated by AlphaFold. The command “color bfactor palette alphafold” was used to color-code the confidence for each prediction (scale from red = 0% confident - blue = 100% confident). Genes encoding proteins in structure clusters 7 and 8 are over-represented in WGCNA M4 and M5.

### Stage-specific expression of Mep effectors of *M. oryzae*

In order to focus on the most likely effector candidates for experimental analysis, we selected 63 *MEP* genes identified in all experiments (Figure 3C and Supplemental Data Sets 3-5) and functionally characterized 32 *MEPs* by live-cell imaging and targeted gene replacement. We first generated GFP fusions of the 32 representative *MEP* genes, expressed these GFP fusions in Guy11, and visualized their sub-cellular localisation patterns during fungal growth *in planta*. A sub-set of effectors could be visualised very early during plant infection. *MEP1*, for example, is organized into a ring conformation at the appressorium pore on the plant cell surface (Figure 5). The appressorium pore is organised by septin GTPases and marks the point of penetration peg emergence and polarised exocytosis, mediated by the exocyst complex (Gupta et al., 2015). Mep1 is therefore likely to be secreted at the point of cuticle penetration from the penetration peg. After cuticle rupture, Mep1-GFP initially accumulates at the tip of primary invasive hypha and then outlines invasive hyphae, suggesting that it localises to the apoplast between the fungal cell wall and the plant plasma membrane, termed the Extra-Invasive Hyphal Membrane (EIHM) (Kankanala et al., 2007), as shown in Figure 5B and Supplemental Movie 2. Consistent with this idea, Mep1-GFP co-localises with Bas4-RFP, an apoplastic effector (Giraldo et al., 2013) around invasive hyphae 24h after infection, consistent with its classification in M4 (Figure 5B). By contrast, the M5 candidate effector gene*, MEP3*, localises exclusively to the BIC during infection, peaking in expression at 48h (Figure 5C). Exclusive BIC localisation was observed for all other Mep-GFP fusions examined within the initial cell penetrated (Supplemental Figure 5).

**Figure 5.**
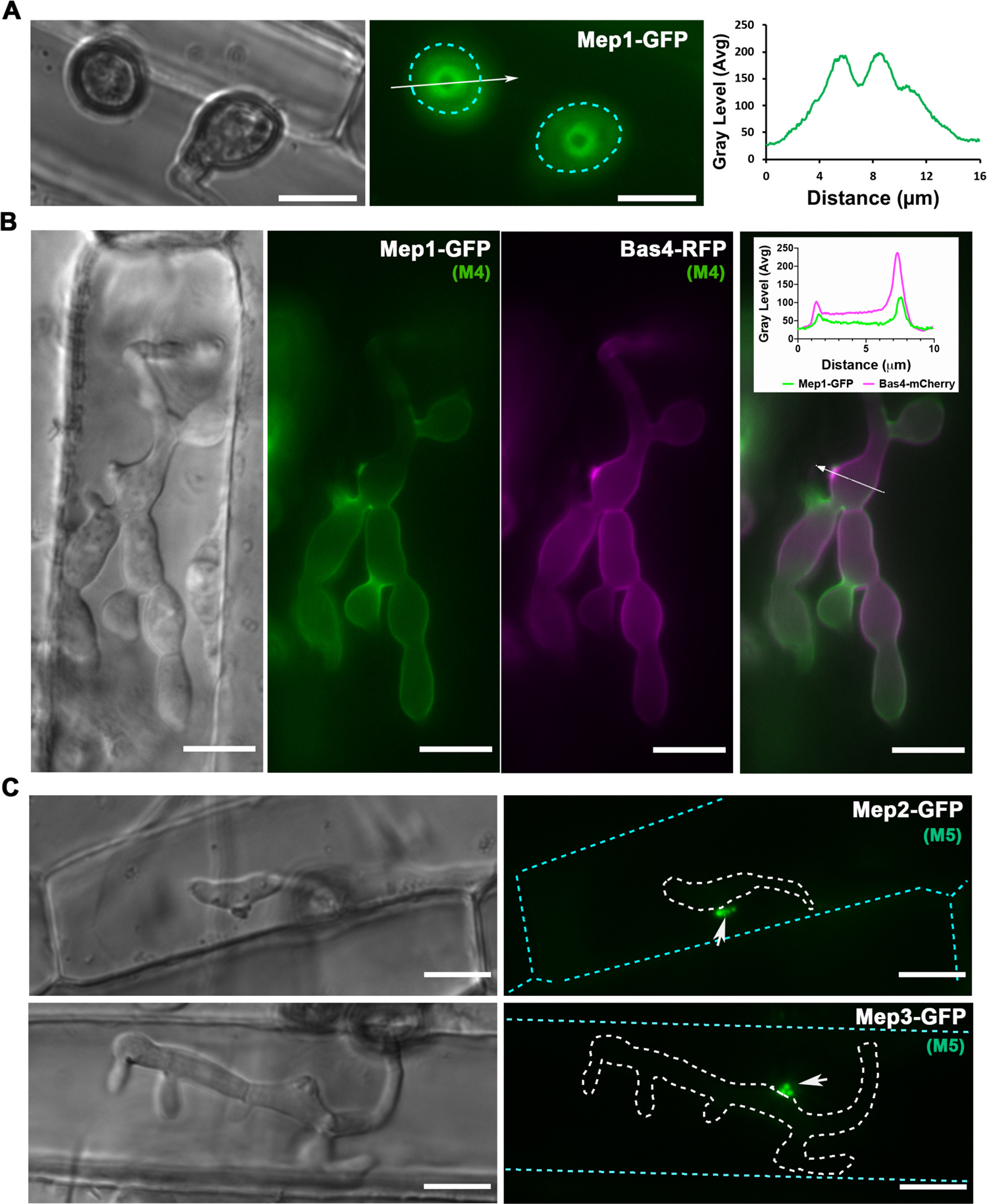
Live-cell imaging of Mep candidates during plant infection reveals spatial localisation of effectors during appressorium penetration and invasive growth. **(A)** Micrographs showing the M4 effector Mep1-GFP localising in a ring confirmation at the appressorium pore at the leaf surface in *M. oryzae* Guy11. Conidial suspensions at 5 ×10^4^ mL^-1^ were inoculated onto rice leaf sheath and images captured at 24 hpi. The periphery of the appressorium is indicated by a cyan dotted line. Line scans show Mep1-GFP fluorescence in a transverse section of the appressorium. Scale bars = 10 µm. **(B)** Conidia were harvested from a *M. oryzae* transformant expressing Mep1-GFP and Bas4-RFP gene fusions and inoculated onto rice leaf sheath preparations. Images were captured at 28 hpi of invasive growth. Micrograph and linescan graphs show co-localization of Mep1-GFP and Bas4-RFP in invasive hyphae of the *M. oryzae* wild-type strain Guy11. Scale bars = 10 µm. **(C)** Micrographs showing localisation of representative M5 effector candidates Mep2-GFP and Mep3-GFP from the 32 selected Mep proteins (Supplemental Figure 9) during plant infection. Images were captured at 22 hpi-24 hpi. Invasive hyphae are outlined by a white dotted line and individual plant cells indicated with a cyan dotted line. Arrow indicates localisation of both Mep2-GFP and Mep3-GFP at the biotrophic interfacial complex (BIC). Scale bars = 10 µm.

To investigate secretion of MEP effectors, we generated alleles in which the predicted signal peptide region was removed and expressed these in *M. oryzae*. When Mep1^19-74^-GFP was expressed with Mep1-mCherry in Guy11, we observed cytoplasmic accumulation of Mep1^19-74^-GFP, while Mep1-mCherry was delivered to the apoplast around invasive hyphae (Figure 6A and Supplemental Figure 11). The signal peptide of cytoplasmic effectors was also necessary to enable delivery to the BIC, with Mep3^25-145^-GFP fluorescence observed intracellularly in invasive hyphae (Figure 6B). Because the punctate localisation of BIC effectors might be interpreted as being a lower expression level than predicted by RNA-seq analysis (Figure 6C), we also made gene fusions expressing free GFP driven by each *MEP* promoter, as shown for *MEP1* and *MEP3* in Figure 6D and E. Fluorescence was observed for *MEP1* in conidia and appressoria, as well as invasive hyphae, while *MEP3* was expressed at a very high level exclusively in invasive hyphae (Figure 6D and E and Supplemental Figure 10). Cell biological visualisation of MEP expression was therefore consistent with predictions from RNA-seq analysis.

**Figure 6.**
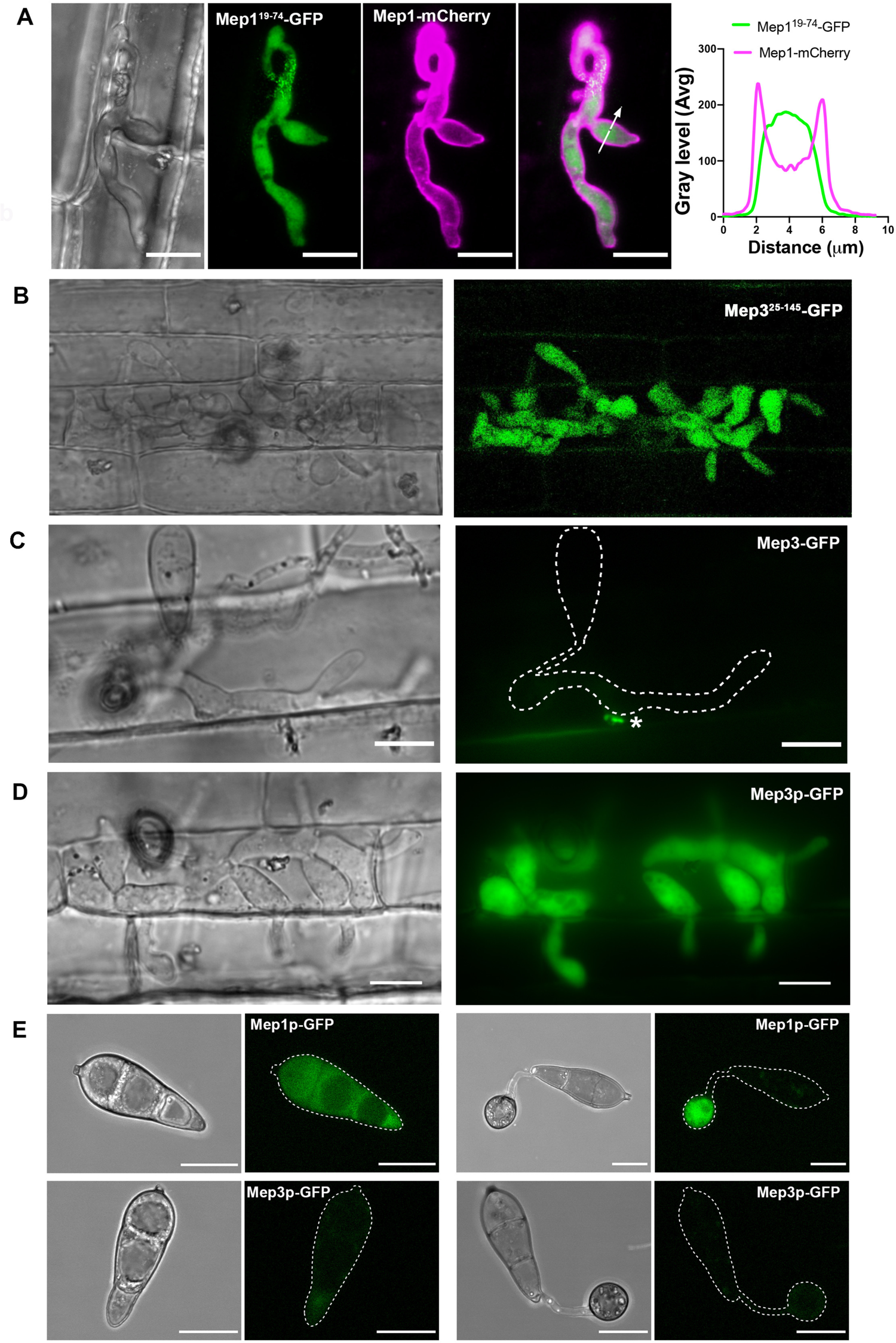
*MEP* genes are highly expressed in living plant tissue during plant infection. **(A)** Conidia were harvested from a *M. oryzae* transformant expressing a Mep1^Δ19-74^-GFP signal peptide deletion and a Mep1-RFP full length gene fusion and inoculated onto CO39 rice leaf sheath. Images were captured at 24 hpi of invasive growth. Micrograph and linescan graphs show that the Mep1 signal peptide is required for correct protein secretion. Scale bars = 10 µm. **(B)** Micrographs showing the rice plant infected by the *M.oryzae* wild-type strain Guy11 expressing a Mep3^Δ25-145^*-*GFP signal peptide deletion. Green fluorescence was retained inside the invasive hyphae, while red fluorescence (false coloured to magenta) was exported to the apoplast. Images were captured at 24 hpi of invasive growth **(C)** Micrographs showing rice plant tissue infected by the *M. oryzae* wild-type strain Guy11 expressing full length *MEP3,* driven by its native promoter. Mep3-GFP could be observed as small puncta at the BIC. Images were captured at 24 hpi of invasive growth **(D)** Micrographs showing rice CO-39 infected with *M. oryzae* wild-type strain Guy11 expressing cytoplasmic GFP driven by the *MEP3* promoter. **(E)** Conidia were harvested from *M. oryzae* transformants expressing GFP driven by the promotor of Mep1p-GFP and Mep3p-GFP, respectively, and inoculated onto hydrophobic glass coverslips. Micrographs of Mep1p-GFP showed fluorescent signal in both the conidium and the appressorium, in contrast to Mep3p-GFP which displayed little signal. Appressorium formation was observed at 20 hpi. Scale bars = 10 µm.

### The dynamics of the host-pathogen interface during plant infection

To investigate the dynamics of the host-pathogen interface during plant infection, we infected a transgenic rice line expressing the plasma membrane marker Lti6b-GFP (Kurup et al., 2005; Mentlak et al., 2012), using a Guy11 strain expressing Mep1-mCherry. The Mep1-mCherry fluorescence outlined invasive hyphae in initially colonised epidermal cell, bounded by the EIHM, which showed Lti6b-GFP fluorescence, consistent with delivery of the effector to the apoplast between the fungal cell wall and EIHM. The EIHM maintained integrity during the early stages of the plant infection (Figure 7A), but membrane integrity was lost once the fungus invaded neighbouring cells, resulting in loss of host cell viability (Figure 7B). The Mep1-mCherry signal accumulated in the first invaded cell suggesting rupture of the plant plasma membrane and closure of plasmodesmata. New BICs were, meanwhile, formed immediately upon entry into neighbouring cells enriched with plant membrane.

**Figure 7.**
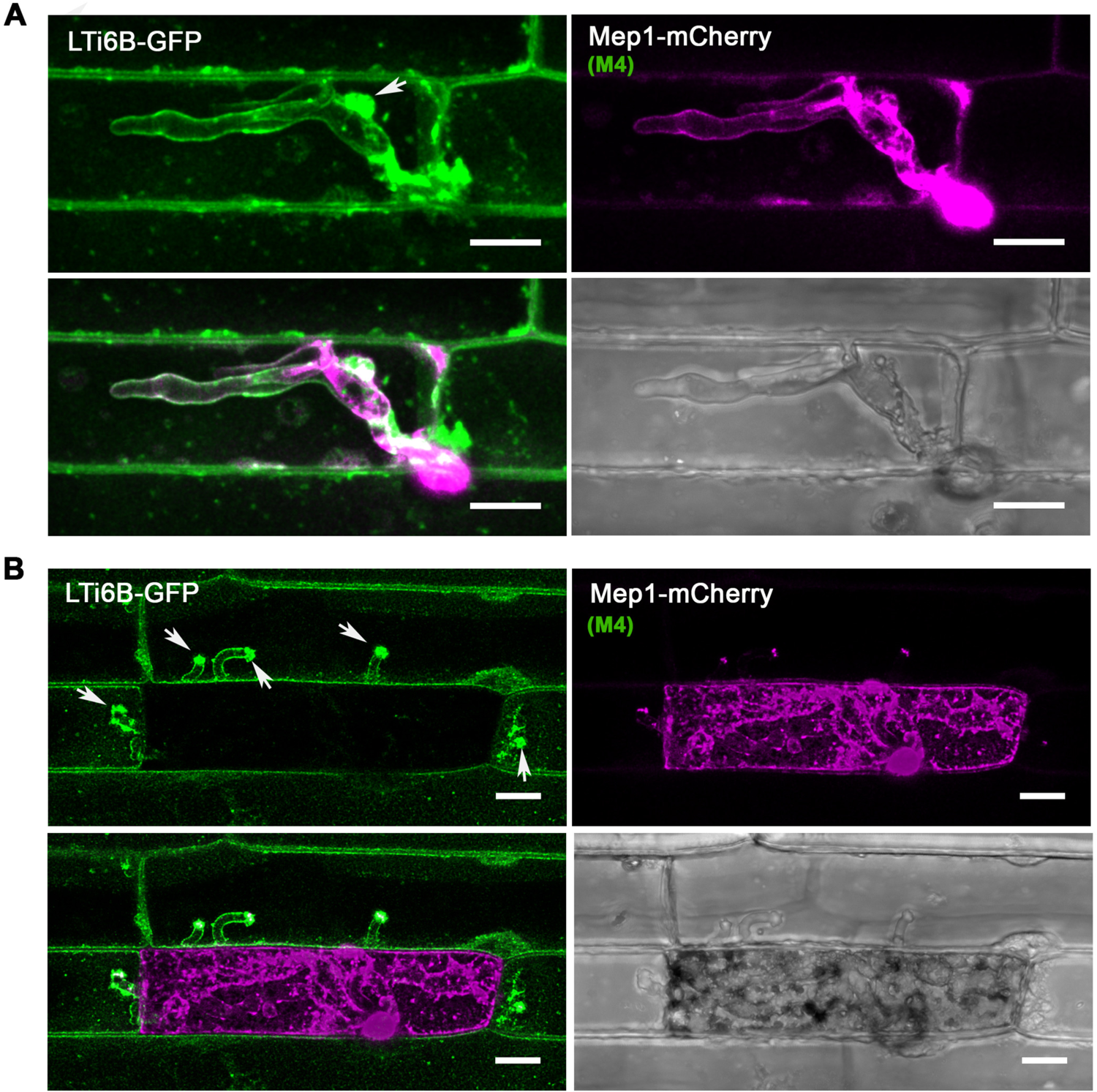
The rice plasma membrane is invaginated and accumulates at the Biotrophic Interfacial Complex (BIC) during plant infection. Laser confocal micrographs of *M. oryzae* expressing Mep1-mCherry colonising epidermal leaf cells of a transgenic rice line expressing plasma membrane-localised LTi6B-GFP. Images were captured at 24 hpi **(A)** and 36 hpi **(B)**. The plant plasma membrane stays intact and invaginated at the early stages of plant infection. LTi6B fluorescence accumulates at the bright biotrophic interfacial complex, indicating the BIC is a plant membrane-rich structure. The fluorescence signal from secreted Mep1-mCherry is surrounded by the fluorescence signal of the rice cell plant plasma membrane marker LTi6B-GFP as the fungus invades new cells, but the initial epidermal cell occupied then loses viability and Mep1-mCherry fluorescence fills the rice cell. Scale bars = 10 um.

### Mep effectors secreted to BICS are always delivered into host cells

Translocation of effectors into rice cells has previously been visualized by expressing fluorescent effector fusion proteins in *M. oryzae* with an artificially added C-terminal nuclear localization signal (NLS) (Khang et al., 2010). The NLS allows effector detection in plant cells by concentrating them in the rice nucleus. The assay has previously been used, for example, to show that the *PWL2* effector, which acts as a host specificity determinant in *M. oryzae,* is translocated into rice cells (Khang et al., 2010). To confirm whether *MEP*-encoded effectors are translocated into host cells, we generated translational fusions with the Mep effector fused in-frame to an NLS. As a control, we expressed GFP-NLS driven by the constitutive TrpC promoter and found that the GFP signal accumulated predominantly in fungal nuclei confirming that GFP-NLS is functional and can drive free GFP into nuclei. However, unless specifically delivered, the signal will not move into a plant nucleus during plant infection (Figure 8A).

**Figure 8.**
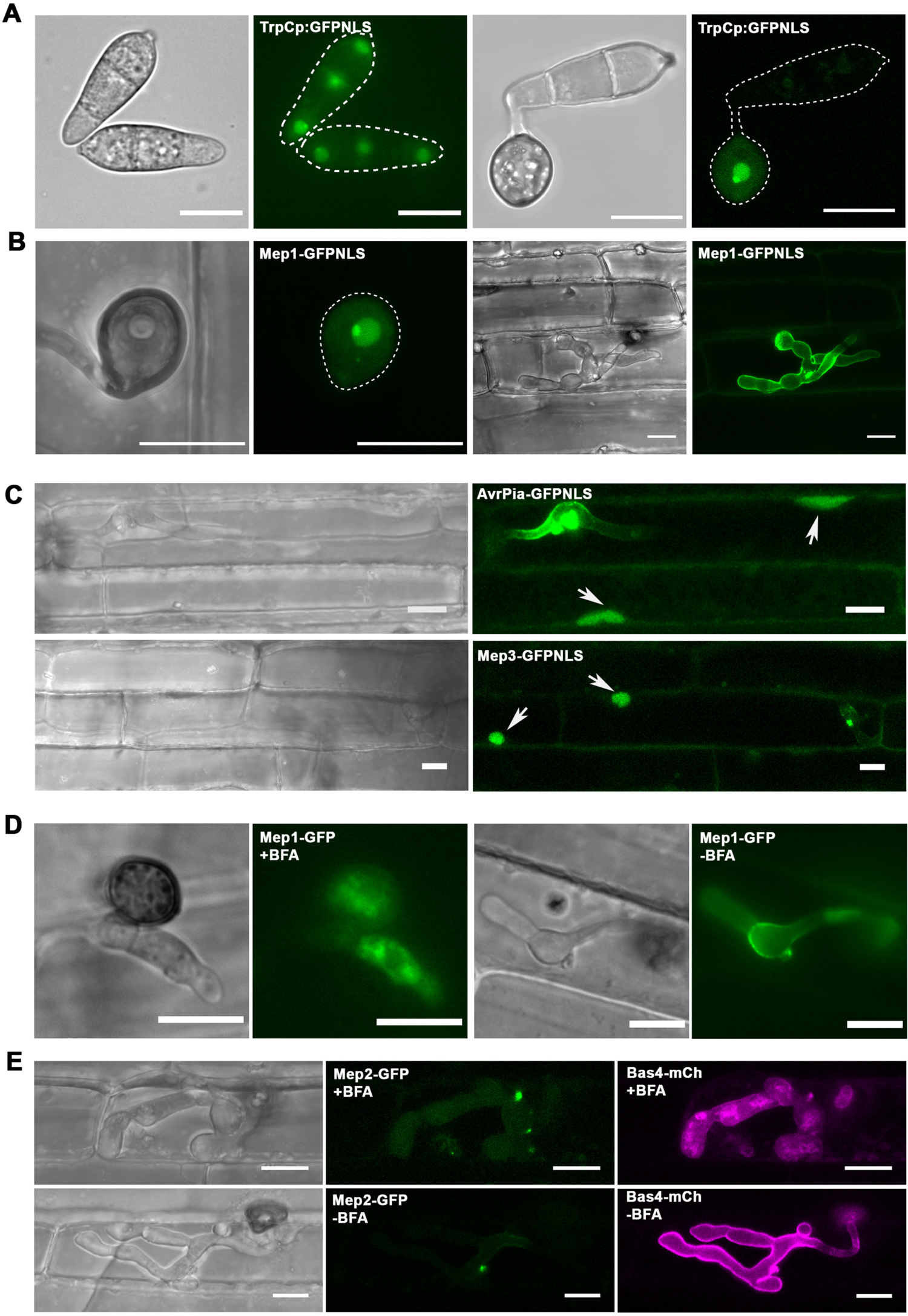
Cytoplasmic Mep candidates, which accumulate at the BIC, are translocated into host cells.**(A)** Cellular localisation of GFP with a nuclear localization signal (NLS) driven by the TrpC promoter in *M. oryzae*. The fluorescence signal in the nuclei and nucleoli of conidia and the appressorium. The outline of the fungus is depicted by a white dotted line. **(B)** Micrographs of Mep1-GFP^NLS^ inoculated onto rice leaf sheath and captured at 28 hpi. Fluorescence could be observed in the nucleus and nucleolus of the appressorium, but not in the fungus or plant cells. **(C)** Micrographs of AvrPia-GFP^NLS^ and BIC-accumulating Mep effector candidates showing delivery into both the invaded host cell and unoccupied neighbouring host cells. Arrows indicate plant nuclei. **(D)** Cellular localisation of Mep1-GFP in *M. oryzae* during biotrophic growth on epidermal rice cells. Brefeldin A (BFA) was applied at 20 hpi and 0.1% DMSO was used for the control treatment. Images were captured at 3-4 h post treatment. Mep1-GFP fluorescence showed accumulation within invasive hyphae and particularly in the BIC-associated cell **(E)** Micrographs of *M. oryzae* expressing Mep2-GFP and Bas4-mCherry during biotrophic growth on epidermal rice cells. BFA treatment was used to look at secretion of Mep2-GFP and Bas4-mCherry. Mep2-GFP fluorescence accumulated in the BIC in the presence or absence of BFA. By contrast, Bas4-mCherry fluorescence accumulated inside invasive hyphae. Scale bars = 10 µm.

To investigate whether the Mep1 effector, which we localised to the apoplast, can be translocated into host cells, we expressed Mep1-GFP-NLS and infected rice seedlings. We found that despite the effector possessing a signal peptide and normally localising to the appressorium pore (shown previously in Figure 4a), fluorescence was observed predominantly in the fungal nucleus (Figure 8B). No GFP signal was observed in the plant nucleus of the occupied rice cell, or surrounding plant nuclei (Figure 8B). By contrast, when the BIC-localised effector Mep3 was fused to an NLS, the Mep3-GFP-NLS signal was observed in the plant nucleus of the occupied cell and those of adjacent cells (Figure 8C). All BIC-localised effectors evaluated using this assay showed translocation to host cells, while apoplastic effectors such as Mep1 were not translocated.

Apoplastic and cytoplasmic effectors of *M. oryzae* have been reported to be secreted by different pathways during rice infection, but to date only a very limited sample has been analysed (Giraldo et al., 2013). Cytoplasmic effectors, such as Pwl2, were shown to be secreted in a brefeldinA (BFA)-insensitive manner, suggesting that they do not undergo conventional ER-to-Golgi secretion. By contrast, apoplastic effectors such as Bas4 are secreted conventionally and are sensitive to BFA treatment. We therefore applied brefeldin A to rice tissue infected with *M. oryzae* expressing Mep-GFP fusions. We observed that secretion of Mep1-GFP was significantly inhibited in the presence of brefeldin A, with fluorescence accumulating within invasive hyphae (Figure 8D). However, when we assayed BIC-associated effectors such as Mep2 and Mep3, we found they were BFA-insensitive. We observed that when a *M. oryzae* Guy11 isolate expressing Mep2-GFP and Bas4-mCherry was exposed to BFA (Figure 8E), then the BAS4-mCherry signal was prevented from being delivered to the apoplast, while the Mep2-GFP signal remained in the BIC. This was consistent for the whole set of analysed putative effectors. We conclude that Mep effectors are secreted by two distinct secretory pathways depending on their host destination (Giraldo et al., 2013).

### Mep effectors contribute to pathogen fitness

Pathogenic fungi secrete a large battery of effector proteins during infection, but individual effector genes have often been reported to display no discernable mutant phenotype with regard to virulence. This has been interpreted as being a consequence of functional redundancy and overlapping effector functions, such that the role of an individual effector is very hard to measure (Selin et al., 2016). In *M. oryzae* most effectors characterized to date– such as Pwl2, Bas1, Bas2, Avr-Pik, and Avr-Pita –for example, have been shown not to contribute substantially to the ability of the fungus to cause disease. A much smaller number of effectors, such as the extracellular LysM effector Slp1, do contribute to fungal virulence (Mentlak et al., 2012) but such reports are very rare. It is, however, obvious from an increasing number of studies that effectors serve important functions in the suppression of plant immunity. They interfere with the operation of pattern recognition receptors, impair host responses such as reactive oxygen species generation, and suppress immunity signaling pathways and transcriptional responses (Irieda et al., 2019; Kim et al., 2020). The contribution of an effector to fungal virulence is normally evaluated by generating a targeted deletion mutant and infecting a susceptible host cultivar. Disease symptoms are then measured compared to an isogenic wild type strain. We decided that this assay was likely to be insufficiently sensitive to be able to accurately determine the contribution of a Mep effector to rice blast disease. We therefore deployed a relative fitness assay (Ross-Gillespie et al., 2007; Lindsay et al., 2016) to evaluate the contribution of an effector to fungal virulence. We generated a Guy11 strain expressing cytoplasmic GFP under control of a high-level constitutive promoter, TrpC. At the same time we generated a null mutant of the specific *MEP* effector gene (Supplemental Figure 12), in this instance *MEP1*, expressing cytoplasmic mCherry driven by the same TrpC promoter. We then generated a spore inoculum in which we mixed wild type GFP spores and *!ιmep1* mCherry spores in a 1:1 ratio. This inoculum was used to inoculate CO39 seedlings. We allowed the disease to progress and then recovered conidia from disease lesions (Figure 9A). We recorded the ratio of spores and then used the same ratio to prepare inoculum to infect a second batch of rice seedlings. We repeated the assay for each generation. We observed that in a mixed infection, the *!ιmep1* mutant was driven close to extinction after three rounds of infection (Figure 9B). Relative fitness was calculated using the equation x_2_(1-x_1_) / x_1_ (1-x_2_), where x_1_ is the initial frequency of the Mep mutant in the population and x_2_ is the final frequency after three generations of infection (Figure 9C). In a control experiment, we mixed the wild type and *!ιmep1* mutant conidia in axenic plate culture and then recovered spores 12 days later. We observed that they maintained the same 1:1 ratio, showing that *!ιmep1* mutants have equivalent fitness when growing outside of a plant (Supplemental Figure 13). In this way, we were able to calculate the fitness cost of losing an individual effector during plant infection and show that Mep1, for example, contributes significantly to rice blast disease.

**Figure 9.**
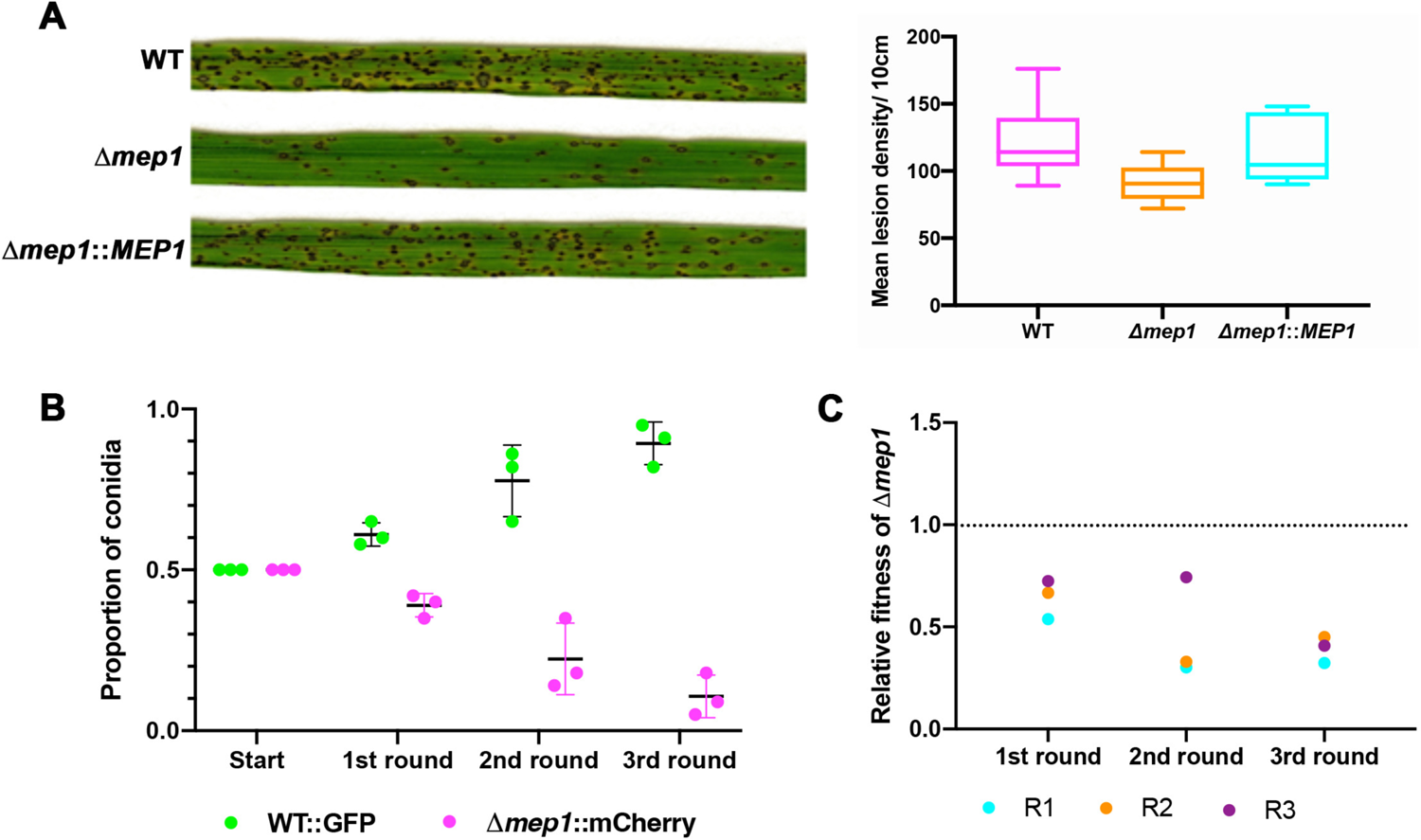
Mep1 contributes to pathogen fitness during plant infection. **(A)** Conidial suspensions of equal concentration (5×10^4^ spores mL^-1^) from *M. oryzae* Guy11 or Δ*mep1* mutants were used to inoculate 21-day-old seedlings of the blast susceptible cultivar CO-39, and disease symptoms recorded after 5 dpi. The box plot shows the mean lesion density of seedlings infected with Guy11 and the Δ*mep1* mutant per unit area. (**B**) A relative fitness assay was carried out by mixing conidia in equal amounts (1:1) from Guy11 expressing GFP and *Δmep1* mutant expressing mCherry. Spores were collected from disease lesions and used to inoculate new seedlings in the recovered proportions. A dot plot showing the proportion of WT::GFP and the Δ*mep1*::mCherry conidia recovered from each generation is shown. The Δ*mep1*::mCherry mutant was driven to near extinction in three generations **(C)** Relative fitness was calculated using the calculation x2(1-x1) / x1 (1-x2), where x1 is the initial frequency of Δ*mep1*::mCherry conidia and x2 the final frequency.

## Discussion

The ability of the rice blast fungus to colonise plant tissue and cause disease is still relatively poorly understood (Fernandez and Orth, 2018; Eseola et al., 2021). The most significant recent advances have come from exploring the cellular changes that accompany fungal infection (Khang et al., 2010; Eseola et al., 2021), the regulation of primary metabolism associated with biotrophic growth (Sun et al., 2018), the secondary metabolic pathways associated with suppression of host immunity (Patkar et al., 2015; Marroquin-Guzman et al., 2017), the definition of effector functions (Mentlak et al., 2012; Kim et al., 2020), and the identification of signalling pathways associated with invasive growth (Sakulkoo et al., 2018). These have provided insight into the substantial changes elicited by *M. oryzae* as it infects rice plants, in order to cause disease (Fernandez et al., 2014; Cruz-Mireles et al., 2021). These advances have been coupled with numerous studies to validate the roles of individual genes in pathogenesis, although these have predominantly been associated with appressorium-mediated infection, with only a small number of gene defined as being determinants of tissue colonisation (Fernandez and Orth, 2018). Our understanding of the biology of rice tissue colonisation has therefore been limited because of a lack of holistic studies of blast infection. The motivation for this study was to carry out transcriptional profiling of the entire disease cycle of *M. oryzae* in order to reveal fundamental changes in pathogen gene expression that occur during progression of the disease, and to use this information to identify the full repertoire of fungal effector proteins deployed by the fungus.

Transcriptional profiling has been used in several plant-pathogen interactions to provide insight into fungal development inside a host plant (Kawahara et al., 2012; Kleemann et al., 2012; Rudd et al., 2015; Dobon et al., 2016; Wang et al., 2017; Lanver et al., 2018). These studies have suggested that expression of effector proteins is an emerging characteristic of fungal invasion of plant hosts. In the corn smut fungus *Ustilago maydis*, transcriptional profile analysis has provided significant insight into the biology of biotrophic development. Distinct temporal expression profiles have, for instance, been defined for a wide range of genes associated with fungal metabolism, nutrient acquisition, regulatory networks and secrete effectors. In particular, *U.maydis* effectors were found to be expressed in three distinct expression modules associated with growth on the leaf surface, biotrophic development in maize cells, and during induction of tumour formation, when rapid plant cell division is stimulated (Lanver et al., 2018). Until now, the studies carried out in *M. oryzae* have not reached such a level of resolution.

The most comprehensive study to date in *M. oryzae* used laser capture microdissection to enhance the proportion of rice cells containing fungal invasive hyphae and then employed microarray analysis to define a total of 58 putative fungal effectors, of which 4 were localised during the infection process (Mosquera et al., 2009). A more recent *M. oryzae* study using RNA-seq analysis, focused on a single time point of infection, 24 hpi, when rice cells were initially occupied by the fungus. This led to 240 fungal transcripts being defined, which included some effector candidates (Kawahara et al., 2012). These studies of *M. oryzae*, although influential, do have severe limitations in their coverage of the development of rice blast and the level of resolution they were able to achieve. Laser capture microdissection, for example, added a layer of perturbation to sample preparation (Mosquera et al., 2009), while focusing on a single time point, (Kawahara et al., 2012) limited the identification of effectors. We therefore decided to perform a comprehensive transcriptional profiling study in which 8 timepoints would be used, along with distinct inoculation methods and different fungal-rice cultivar interactions, to identify the maximum number of fungal transcripts and provide the closest relationship to natural blast infections.

The major findings of our study are (1), the definition of 10 distinct modules of differentially regulated *M. oryzae* genes that encompass the entire process of pathogenic development from the time of spore germination to the development of sporulating disease lesions; (2), the identification of distinct physiological processes and genes involved in secondary metabolism expressed at specific stages of pathogen development; (3), the identification of novel *MEP* effector genes that are temporally expressed throughout pathogenesis in distinct temporal groups; (4), the discovery that structurally conserved effectors are temporally co-regulated during invasive growth, providing a means of identifying novel effector functions; (5), the classification of Mep effectors into cytoplasmic and apoplastic proteins, delivered via two distinct secretory routes, validating this classification system for the effector repertoire, and (6), the definition of Mep effector function based on contribution to pathogen fitness using a mixed inoculation assay and fitness measure.

Temporal analysis of gene expression revealed the nature of changes in physiological function during pathogenic development. This highlighted the rapid growth of *M. oryzae* during spore germination and starvation stress associated with early development on the leaf surface. Consistent with this, genes associated with autophagy, regulated proteolysis, and lipid metabolism were expressed. During early biotrophic colonisation of leaf tissue significantly up-regulated genes associated with carbon and nitrogen source acquisition from the plant host by 24h. The orchestration of secondary metabolic pathways are also a feature of the biotrophic colonisation of initially occupied epidermal cells, in addition to effector secretion. Another key feature recognised during infection is the pattern of repression of gene expression that accompanies the developmental programme following appressorium development. This is consistent with embryogenic specialisation in cell fate, constraining patterns of gene expression and may be a largely unexplored feature of pathogenesis, as so many studies have instead focused exclusively on genes induced during infection.

A pattern therefore emerges of how *M. oryzae* switches from a nutrient-free environment of the leaf surface where its metabolism is dominated by lipid metabolism and re-cycling of the spore contents, consistent with rapid generation of compatible solutes, such as glycerol, for appressorium turgor generation. As the fungus encounters sugars and amino acids within the leaf, it switches to rapidly acquiring nutrients, as it moves through rice cells. A transition to sucrose utilization and the pentose phosphate pathway is apparent, consistent with experimental studies in which the *M. oryzae* glucose-6-phosphate sensor, Tps1, has been shown to regulate carbon metabolism and link glucose availability to glutathione-dependent antioxidation and the establishment of biotrophy (Wilson et al., 2010; Fernandez et al., 2012). The biotrophy-associated gene *IMP1*, for example, which is linked to TOR-dependent nutritional control of invasive growth (Sun et al., 2018) is in M6 and peaks in expression at 48h after infection. Primary metabolism-associated gene expression is also consistent with previous metabolomic analysis of plant infection by *M. oryzae* (Parker et al., 2009), which revealed fungal re-programming of plant metabolism during infection, including suppression of the defence-associated ROS burst. The nitronate mono-oxygenase gene, *NMO2*, for example, which is required to limit nitro-oxidative stress during rice immune responses (Marroquin-Guzman et al., 2017) peaks in expression at 16h and 48h, during the most active phases of biotrophic proliferation of the fungus. Our study also provides evidence of a very large-scale change in gene expression during the later stages of infection after 48h associated with the utilisation of completely distinct families of transcriptional regulators as the fungus transitions to necrotrophy and disease symptom development.

Arguably, the most significant finding of this study is that the effector repertoire of *M. oryzae* is likely to be much more substantial than previously thought, with at least 546 putative effectors recognised, among a total of 863 differentially-regulated secreted proteins. Effector functions are continually being described across phytopathogenic fungi, and in *M. oryzae* a range of effector targets have been identified, including small heavy metal associated domain proteins, inhibitors of chitin-triggered immunity and exocyst components (Franceschetti et al., 2017). In all cases so far described, the effector exerts an effect to suppress pattern-triggered immunity (PTI). However, the sample size evaluated so far is extremely small compared to the likely total repertoire revealed from this study (less than 2%), suggesting that the fungus has the capacity to overwhelm plant defense, perhaps by intervention at key points, with the same targets identified by many effectors, or conversely, suggesting that there are many new effector functions to be discovered, including potentially some not associated with PTI suppression, perhaps inducing morphological changes in rice cells essential for pathogen development.

A limitation in fungal effector identification has been the lack of sequence similarity among effectors and the absence of specific motifs, such as the RXLR motif found in many oomycete effectors (Morgan and Kamoun, 2007). The use of genome-wide computational structural biology approaches does, however, provide a powerful new means of identifying putative effectors, given the structural similarity exhibited by the MAX effectors and ARTs (Seong and Krasileva, 2021a). We observed that both of these structurally conserved effector families are temporally co-expressed within M4 and M5, making them the most enriched for computationally predicted fungal effectors. This has provided a wealth of noew potential effector candidates for functional analysis and also suggests that co-expression in these tow WCGNA modules may provide a new means of mining for novel effector functions, utilizing the combination of structural prediction and temporal expression profiling. It may also be possible to combine these analyses with an evaluation of the chromosomal distribution of gene loci as an additional diagnostic tool for predicting likely effector candidates.

The study of effectors has hitherto also been constrained by the fact that their contribution to the biology of the pathogen has been very hard to assess. This is largely a consequence of the relatively crude assays used to define their role in pathogenesis. Carrying out targeted deletion and simply assaying their ability to generate disease symptoms has not, generally, led to discernible mutant phenotypes. We therefore devised a mixed infection assay designed to allow the relative contribution of an effector to pathogen fitness to be measured. We reasoned that, if an effector makes a small contribution to the fitness of *M. oryzae* during disease development then a mutant lacking that effector will be less likely to survive within a population. We therefore marked strains with different fluorescent markers and produced inocula with equal numbers of spores to infect plants. This provided a simple means of assessing fitness over several generations. In the case of the *MEP1* effector gene, targeted deletion resulted in only a small change in disease lesion generation, but in mixed infection the */′ιmep1* mutant was driven to extinction in three generations. This simple assay has considerable utility for evaluating effector function in future and could be further refined by bar-coding to enable mixed populations to be defined in greater detail and with numerous effector mutants within a given population.

In summary, this study has provided a resource for researchers to ask specific questions regarding the major transcriptional changes associated with rice blast disease, including both repression and activation of gene functions. The analyses reported here are a very small element of what could be addressed with the data sets, which offer the deepest and most extensive transcriptome data available for the rice-*M. oryzae* interaction. Furthermore, we have identified a very large battery of more than 546 *M. oryzae* effector candidates and classified these by temporal expression profile and structural conservation, enabling new insights into the range of effector functions deployed by the fungus to cause rice blast disease.

## Materials and Methods

### Fungal and plant growth conditions

Fungal strains and rice plant cultivars were maintained, as described previously (Talbot et al., 1993b). Ten-day old plate cultures of *M. oryzae* were used to collect conidia for appressorium development assays, leaf sheath infections and pathogenicity assays. Infected rice plants were incubated in a chamber at 24 °C, 12 h photoperiod and 90 % relative humidity.

### Preparation of healthy and infected rice samples for RNA-Seq analysis

*M. oryzae* conidia were harvested from 10-day old complete medium (CM) plates (Oses-Ruiz et al., 2021). To carry out plant infections by spray assay, 21-day-old, 3-leaf-stage rice plants were sprayed with conidia of Guy11 at 1×10^5^ spores mL^-1^ in 0.25% (w/v) gelatin. Infected rice leaves were harvested at 8, 16, 24, 48, 72, 96 and 144 hours post inoculation. Rice leaves sprayed with 0.25% gelatin were used as controls to compare differentially expressed plant genes. To carry out plant infections by leaf drop infection, the third leaf of 3-leaf-stage rice plant was placed on a flat surface and inoculated with a 20µL suspension of 1×10^6^ spores mL^-1^ in 0.25% gelatin. A total of 20 lesions were harvested at each time point (8, 16, 24, 48, 72, 96 and 144 hours post inoculation). Rice leaves inoculated with 0.25% gelatin were used in control experiments to compare differentially expressed plant genes. All collected samples were frozen in liquid nitrogen immediately for RNA extraction. Each experiment has 3 biological replicates.

### Total RNA purification and RNA sequencing

Total RNA from each sample was extracted using RNeasy Plant Mini Kit (QIAGEN). RNase-Free DNase Set (QIAGEN) was used to remove all genomic DNA. RNA quantity and integrity were measured by Agilent 2100 Bioanalyzer, and purified RNA was used to make sequence libraries using True-Seq RNA sample preparation kit from Illumina (Agilent). Sequencing was carried out using HiSeq 2500 standard mode. The raw data have been deposited to ENA with accession number PRJEB45007.

### Differential gene expression analysis

Raw reads were separated by Taxonomy ID of *M. oryzae* and *O. sativa* (NCBI: txid 318829 and txid 4530) using Kraken 2 (Wood et al., 2019). The extracted fungal reads from Kraken 2 were used to quantify the abundance of transcripts using Kallisto (Bray et al., 2016). TPM values were used to perform fungal mass estimation, PCA analysis, and an overview of expression profiling analysis. Differential gene expression analysis was performed using Sleuth (Pimentel et al., 2017). Genes defined as up-regulated by having log2 fold change >1 and P-adjust < 0.05. Amino acid sequences of all coding genes from *M. oryzae* 70-15 were used to predict effectors. Venn diagram was generated using jvenn to compare different datasets (Bardou et al., 2014).

### Co-expression analysis and Pathway enrichment analysis

WGCNA was used to analyse the gene co-expression network (Langfelder and Horvath, 2008). Only genes with at least 5 fungal reads from all three replicates were considered. A total of 11990 genes were therefore processed by WGCNA (version 1.69). The function Blockwisemodules was used to produce a network of a Pearson correlation matrix to exam similarity between genes. A Soft power threshold of 12 was chosen because it was the lowest power to obtain the lowest correlation value (0.85) from topology analysis. Module detection was generated by modified settings to minimize numbers of clusters by using minModulesSize=100, mergeCutHeight=0.30. For each module, the expression level of the module eigengene was calculated to visualize co-expression patterns. For KEGG enrichment analysis, clusterProfiler (version 3.11) was used to perform Benjamini-Hochberg tests to gain P-values and q-values(Yu et al., 2012). The enriched metabolic pathways were selected with a p-value < 0.05.

### Construction of vectors and transformation of *M. oryzae*

All 32 selected effector candidates were cloned with their native promoters and entire protein coding sequences without stop codons, then tagged at the C-terminus with GFP using recombination *in vivo* in yeast (Oldenburg et al., 1997). In brief, the linearized pNEB-Nat-Yeast1284 cloning vector was used which contains the *URA3* gene, allowing complementation of uracil (-) auxotrophy. The positive in-fusion plasmid was transformed into *E. coli* to obtain plasmid DNA for fungal transformation. To construct Mep1-GFP-NLS, primers TrpC-F-SpeI and TrpCp-R-EcoRI were used to amplify the TrpC promoter fragment from the hygromycin resistance gene cassette. Primers GFP-F-EcoRI and 3xNLS-R were used to obtain GFP-3xNLS from Addgene plasmid pEGFP-C1 EGFP-3xNLS. Primers TtrpC-nls-F and GFP-KpnI-R were used to amplify the TrpC terminator. All amplified fragments were inserted into linearized pCB1532 digested with SpeI and KpnI to generate TrpC-GFP-3xNLS-TtrpC. To generate Mep1-GFP-NLS, TrpC-GFP-3xNLS-TtrpC was digested with SpeI and EcoRI and the in-fusion clone containing the Mep1 promoter and coding sequence, amplified with primers 353ProNLS-F and 353NLS-R from Guy11 genomic DNA. Confirmed positive vectors were transformed into protoplasts of corresponding *M. oryzae* strains, as described previously (Talbot et al., 1993a).

### Live-cell imaging of *M. oryzae* during vegetative and invasive growth

For imaging conidia and mycelium of *M.oryzae*, samples were incubated on glass coverslips mounted with water. For imaging appressorium development, conidia were placed on inductive hydrophobic glass coverslips and samples checked during appressorium development. To visualize spatial localisation of *M.oryzae* proteins of interest during plant infection, conidia were inoculated into 3-4 week old leaf sheath samples of rice cultivar CO-39.and infections allowed to proceed for 0h-48h. Infected leaf sheaths were trimmed before microscopy (Kankanala et al., 2007). Confocal microscopy was carried out using a Leica SP8 laser confocal microscope. Excitation/emission wavelengths were 488 nm/ 500-530 nm for eGFP, and 561 nm/ 590-640 nm for mCherry. Images were analysed using Leica software and ImageJ. An illustration of key stages during rice blast disease development in Figure 2B was created with BioRender (https://biorender.com/).

### Statistical analyses

Statistical analyses were performed using the software Prism8 GraphPad. Data sets were tested for normal distribution before comparison. They were analysed using unpaired two-tailed Student’s t-test with Welch’s correction when data sets were normally distributed. When data sets were non-normally distributed, the non-parametric Mann-Whitney test was used for comparisons.

## Supporting information

Supplemental Data Set 1

Supplemental Data Set 2

Supplemental Data Set 3

Supplemental Data Set 4

Supplemental Data Set 5

Supplemental Data Set 6

Supplemental Data Set 7

Supplemental Data Set 8

Supplemental Data Set 9

Supplemental Data Set 10

Supplemental Data Set 11

## Acknowledgements

We acknowledge assistance from Dr D.M. Soanes and Dr M. J. Kershaw from the University of Exeter at the beginning of this project. We are grateful for the discussion with F. Menke, Y. Gupta, R. K. Shrestha, J. Win and S. Kamoun from The Sainsbury Laboratory. This work was supported by The European Research Council under the European Union’s Seventh Framework Programme (FP7/2007-2013)/ERC grant agreement no. 294702 GENBLAST and by an award from The Gatsby Charitable Foundation to NJT.

## Author contributions

X.Y. and N.J.T. conceived and designed the project. X.Y., B.T., L.S.R., V.M.W., A.B.E., N.C., A.J.F and M.O. performed experimental work. X.Y., B.T. and D.M performed bioinformatic analysis. X.Y. and N.J.T. wrote the paper with help and input from co-authors.

Correspondence and requests for materials should be addressed to N.J.T.

## Supplemental Data

**Supplemental Figure 1.**
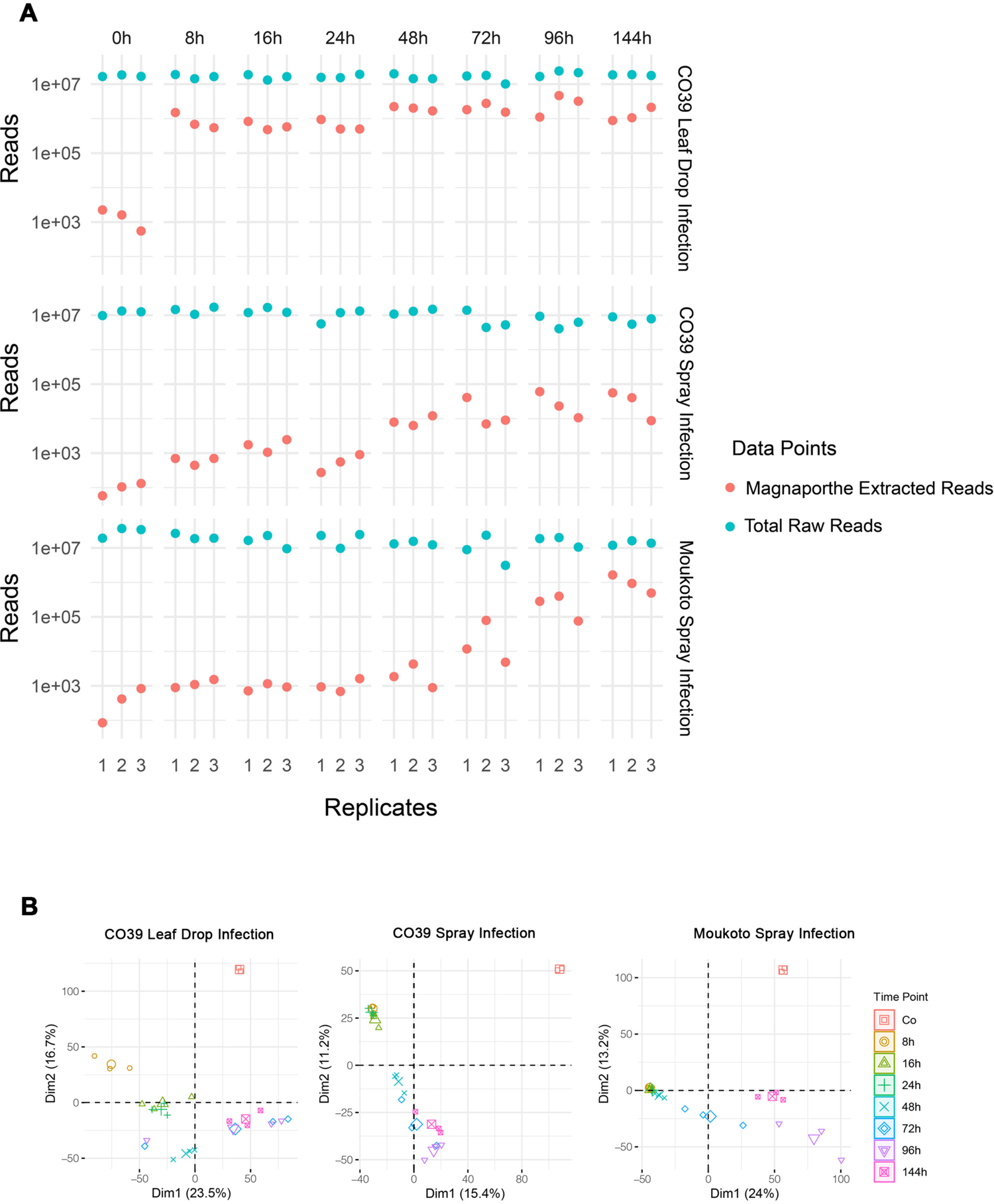
Assessment of the RNA-Seq Data Set of *M.oryzae* during rice infection. **(A)** Graph showing comparison of total raw reads and *M. oryzae* extracted reads from each inoculation method and cultivar-strain combination. **(B)** Principal Component Analysis (PCA) of fungal reads from the three infection datasets; CO39 Leaf Drop Infection, CO39 Spray Infection and Moukoto Spray Infection. Three independent biological replicates were generated for each time point 0h, 8h 16h, 24h, 48h, 72h, 96h, 144h.

**Supplemental Figure 2.**
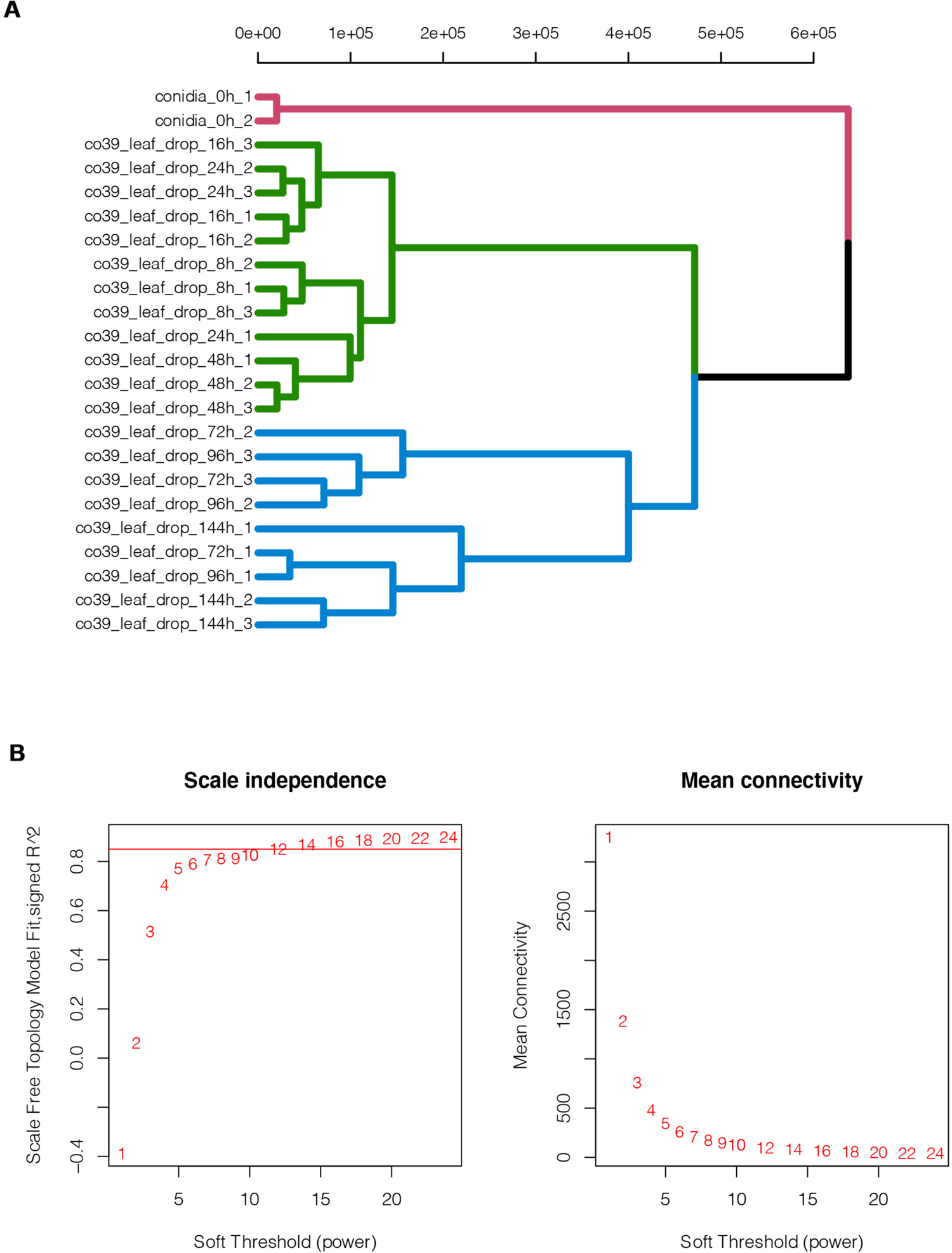
Sample clustering and the soft power analysis for WGCNA analysis. **(A)** Sample cluster analysis of all infected rice RNA-seq data sets. Conidia RNA-seq data sets are significantly different to those from the infection time course (shown in red). RNA-Seq Data Sets from early time points of infection, 8h, 16h, 24h and 48h, all clustered into a single clade (shown in green) while later time points from 72h, 96h and 144h grouped together into a third clade (shown in blue). **(B)** Soft power analysis to obtain the optimised power number for WGCNA analysis. The cut-off threshold indicated that the total RNA-Seq Data Set from infected rice samples could be divided into 10 co-expression modules.

**Supplemental Figure 3.**
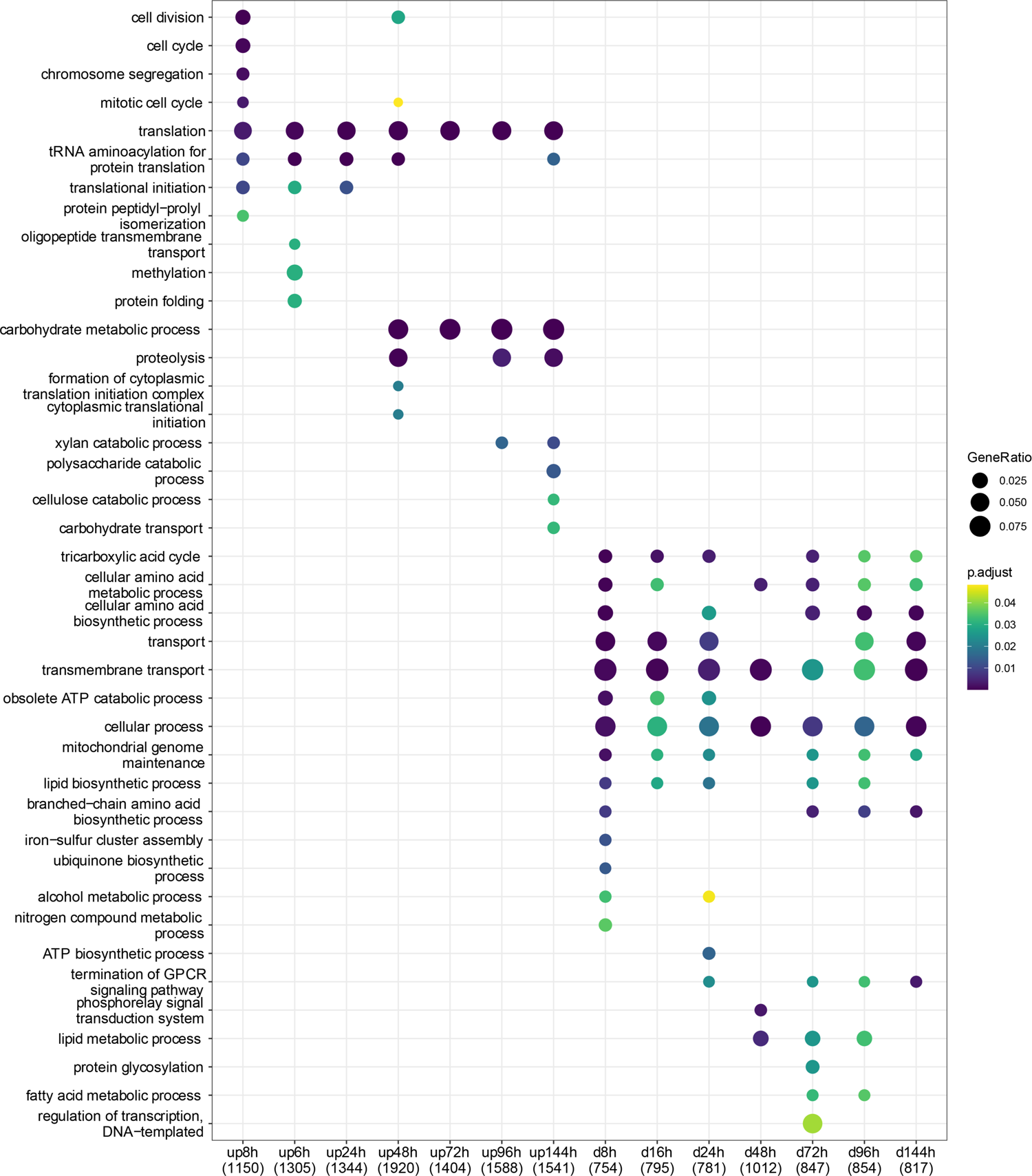
Biological process enrichment analysis of *M. oryzae* genes differentially expressed during plant infection. *M. oryzae* differential gene expression during plant infection was evaluated by comparison to expression in conidial mRNA using Sleuth (Pimentel et al., 2017), identifying genes showing a log2 fold change > 1, and P-adj < 0.05. Diagram shows up-regulated gene functions (top and marked ‘up’ in legend) and down-regulated gene functions (bottom and marked ‘d’ in legend) during a time series of *M. oryzae* infection based on leaf drop inoculation of rice cultivar CO-39 by strain Guy11. GO enrichment analysis was carried out using the R package “mogo” (https://github.com/TeamMacLean/mogo).

**Supplemental Figure 4.**
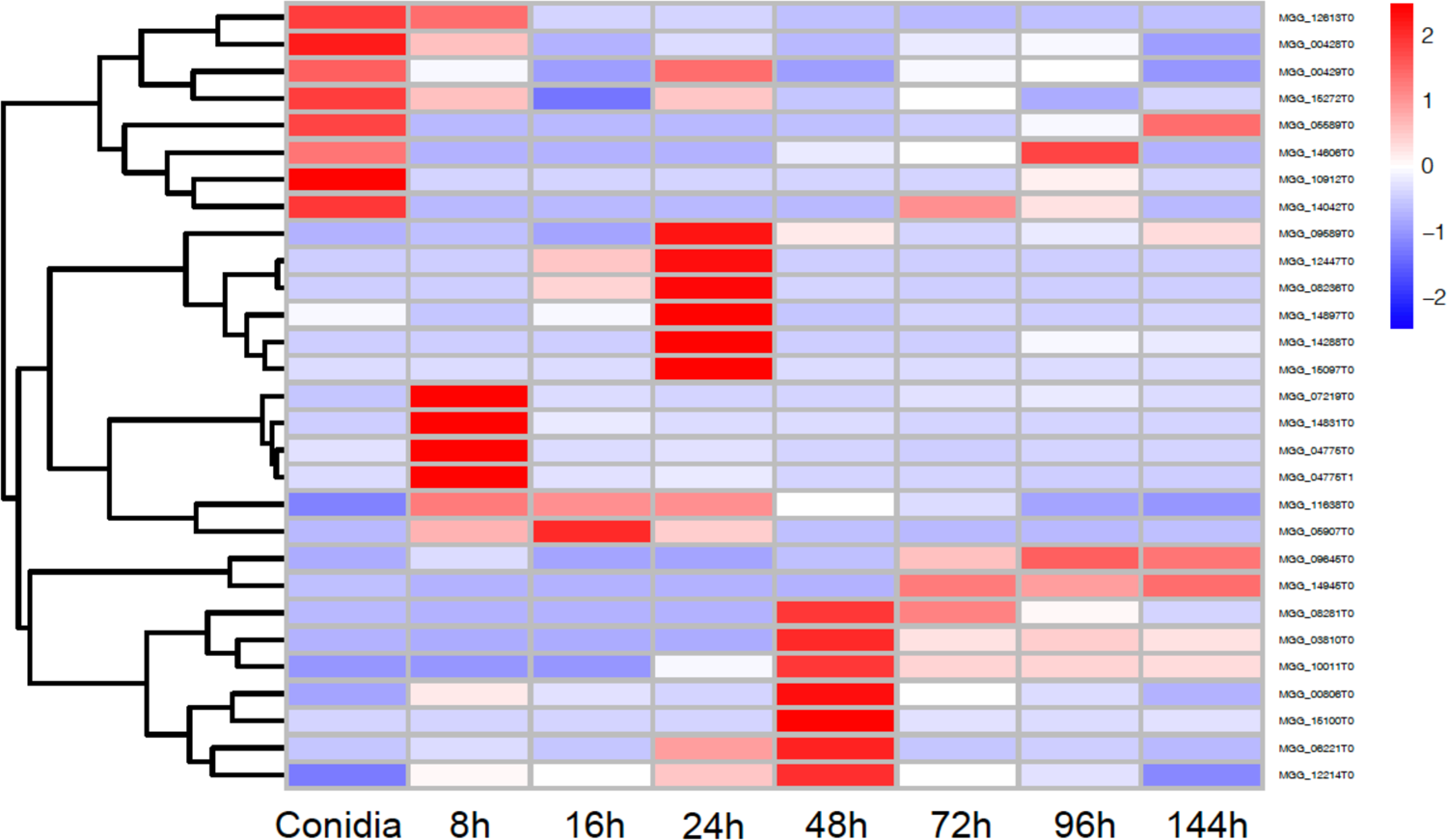
Hierarchical clustering of the expression of *M. oryzae* genes predicted to encode polyketide synthase during plant infection. Heat map showing the temporal pattern of the relative transcript abundance of 29 genes predicted to encode polyketide synthase during plant infection. Genome accession numbers are provided for each gene. Each gene is scaled to the mean TPM (transcript per kilobase million) value across all stages of infection-related-development and plant infection, with fold differences indicated according to the key.

**Supplemental Figure 5.**
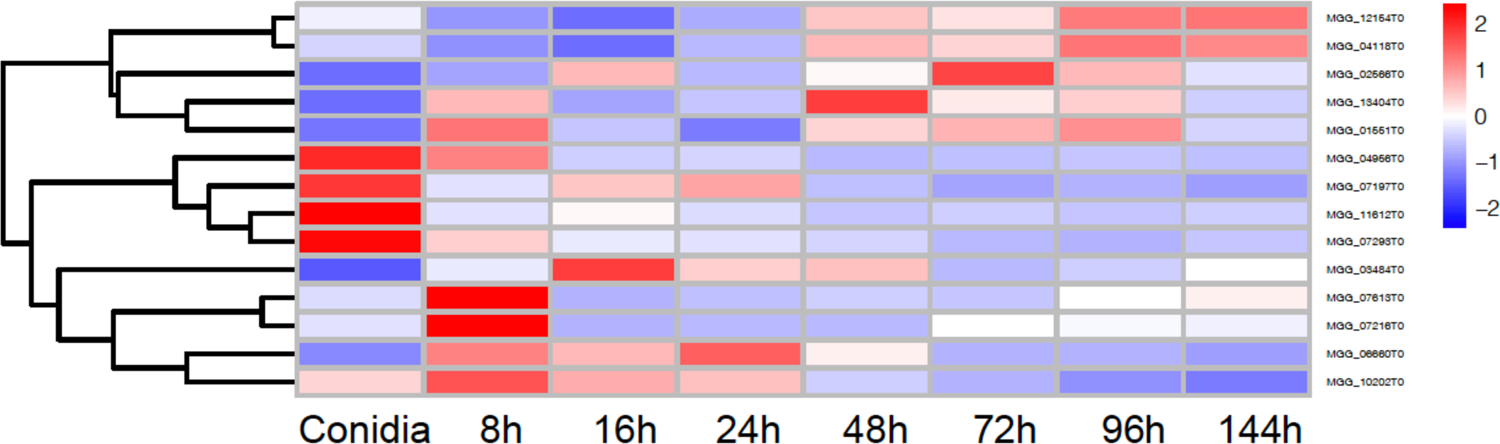
Hierarchical clustering of *M. oryzae* genes predicted to encode fatty acid synthases during plant infection. Heat map showing the temporal pattern of the relative transcript abundance of 14 genes predicted to encode fatty acid synthases during rice infection. TPM (transcript per kilobase million) value across all stages of infection-related-development and plant infection, with fold differences indicated according to the key.

**Supplemental Figure 6.**
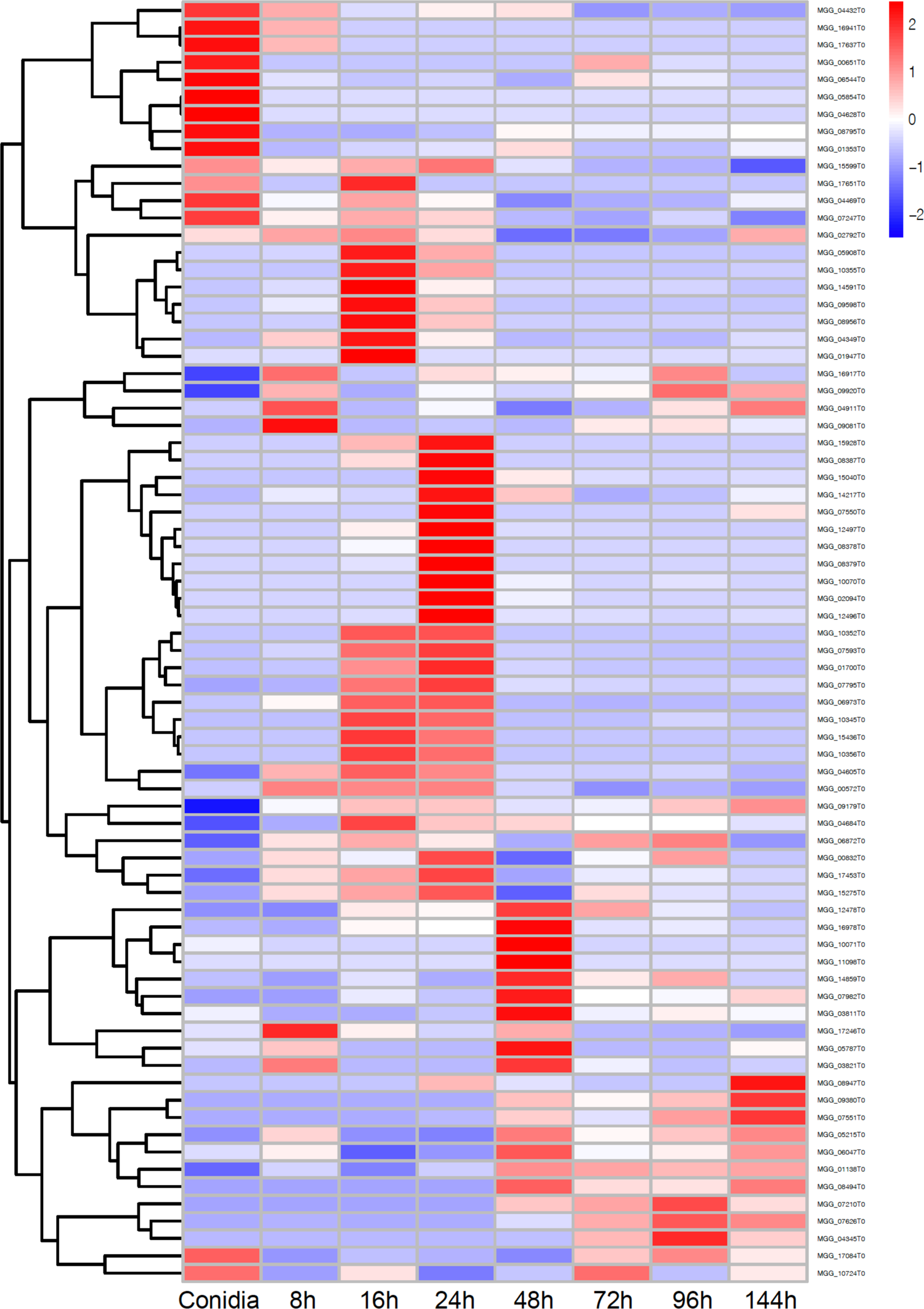
Hierarchical clustering of *M. oryzae* genes predicted to encode cytochrome P450 mono-oxygenases during plant infection. Heat map showing the temporal pattern of the relative transcript abundance of 75 genes encoding cytochrome P450 mono-oxygenases during rice infection. TPM (transcript per kilobase million) value across all stages of infection-related-development and plant infection, with fold differences indicated according to the key.

**Supplemental Figure 7.**
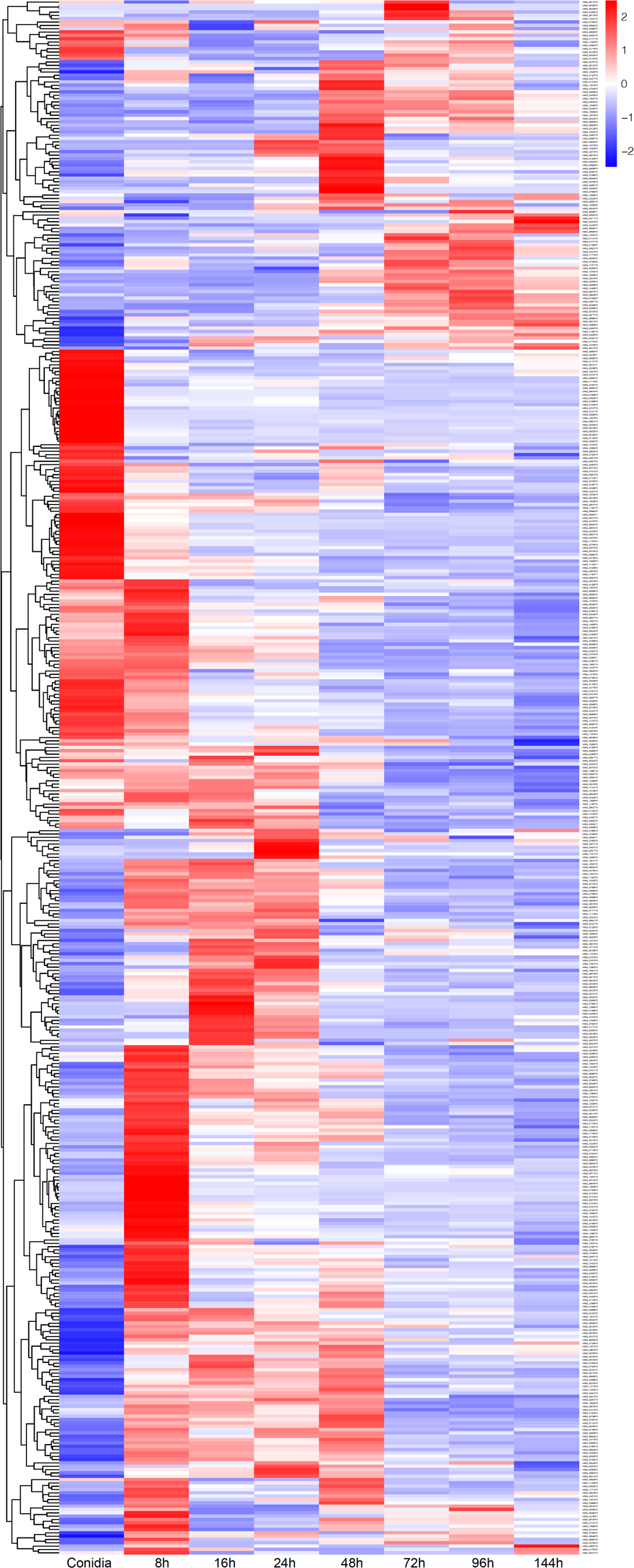
Hierarchical clustering of *M. oryzae* genes predicted to encode transcription factors during plant infection. Heat map showing the temporal pattern of the relative transcript abundance of 495 genes predicted to encode transcription factors during rice infection. All putative transcription factors were extracted from the Fungal Transcription Factor Database (http://ftfd.snu.ac.kr/magnaporthe), a platform designed to identify transcription factor-encoding genes in fungi. TPM (transcript per kilobase million) value across all stages of infection-related-development and plant infection, with fold differences indicated according to the key.

**Supplemental Figure 8.**
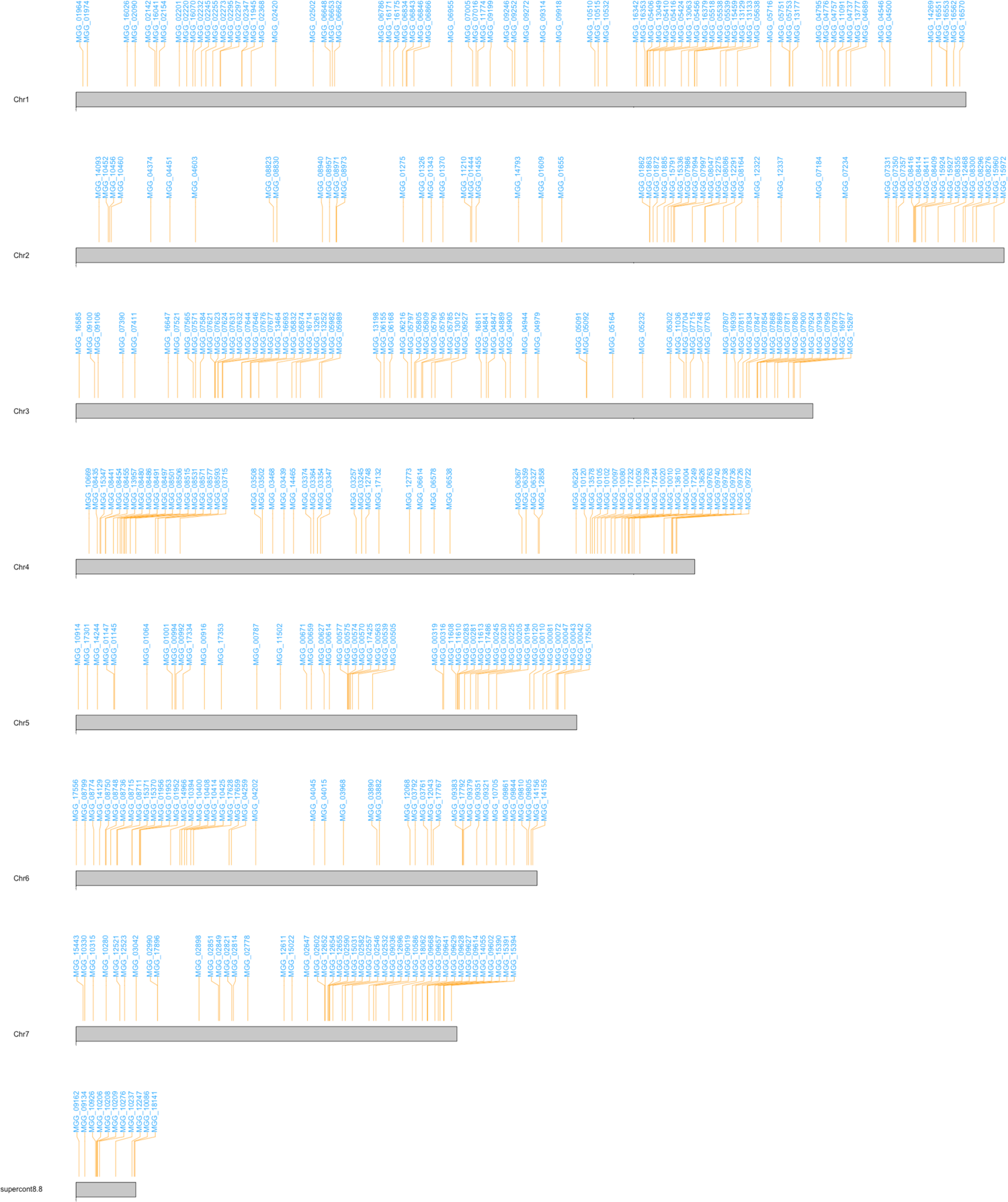
Visualisation of the distribution of 863 *MEP* gene loci on the seven chromosomes of *M. oryzae*. Genetic map showing the order and relative distribution of *MEP* loci among the seven chromosomes of *M. oryzae*. The positions were determined by analysis against the 70-15 reference genome (Dean et al., 2005).

**Supplemental Figure 9.**
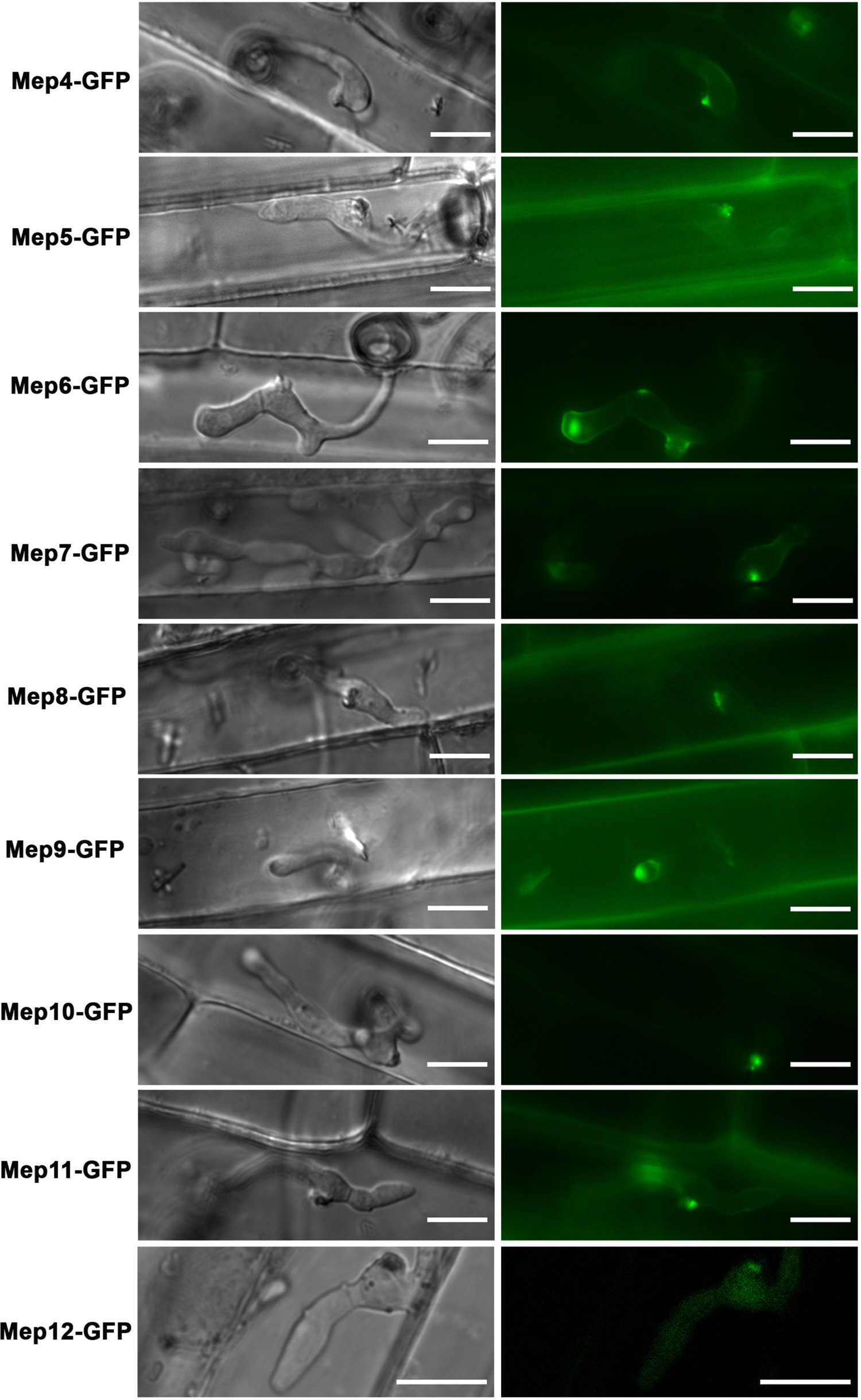
Cytoplasmically targeted Meps consistently localise to the BIC during biotrophic invasive growth of *M. oryzae*. Micrographs showing the localisation of Mep4, Mep5, Mep6, Mep7, Mep8, Mep9, Mep10, Mep11 and Mep12 which were all C-terminally tagged with GFP, transformed and expressed in Guy11 and used to carry out infections of rice leaf sheath of cultivar CO39. Laser confocal images were captured at 22-28 hpi. A single large fluorescent punctum, the BIC, was always observed in the initially invaded epidermal cell. Scale bars = 10 µm.

**Supplemental Figure 10.**
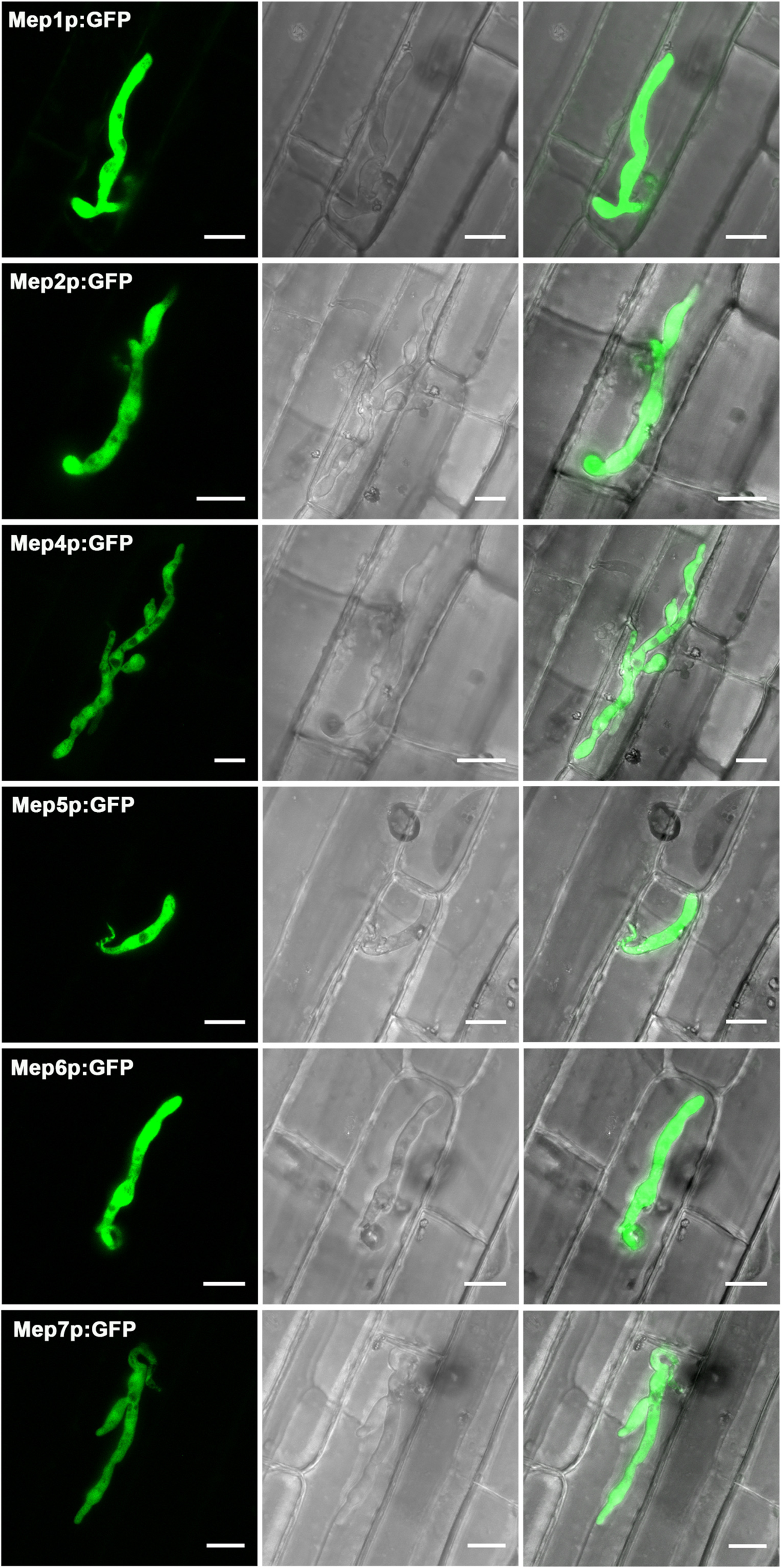
Assessment of the expression of Mep effector candidates during plant infection. Micrographs showing the expression of cytoplasmic GFP driven by the promoters of Mep effector candidates during plant infection. Promoters of *MEP1, MEP2, MEP4, MEP5, MEP6* and *MEP7* were fused to GFP, transformed into Guy11 and single copy transformants selected. A strong fluorescence signal was specifically observed inside invasive hyphae during plant infection. Scale bars = 10 µm.

**Supplemental Figure 11.**
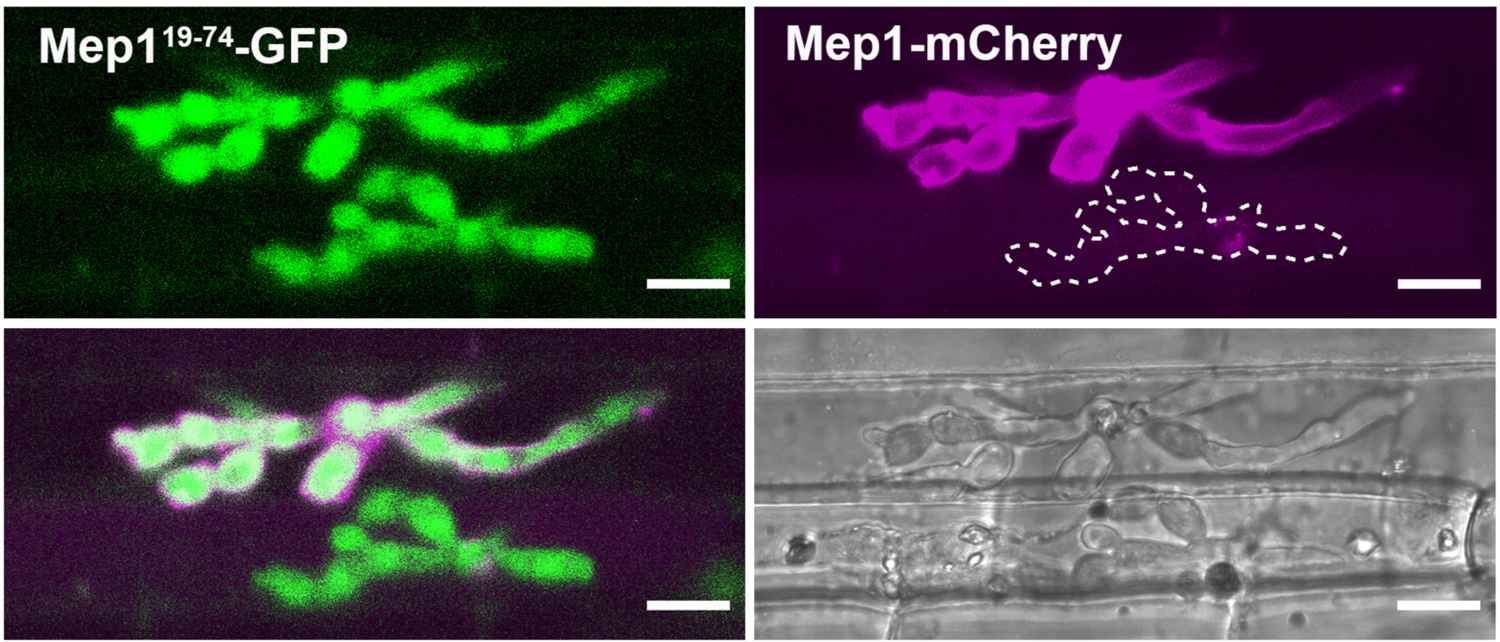
Live-cell imaging of the secreted and non-secreted Mep1 protein variants during plant infection. Micrographs showing invasive hyphae at two infection sites of *M. oryzae* Guy11 expressing Mep1^19-74^-GFP and Mep1-mCherry. Two infection sites are present and can be observed by the GFP fluorescence signal from Mep1^19-74^-GFP and by observation in the bright field channel. At one infection site, the green fluorescence from Mep1^19-74^-GFP is uniformly enveloped by the magenta fluorescence from Mep1-mCherry. This is consistent with delivery of Mep1-mCherry into the apoplast between the fungal cell wall and extra-invasive hyphal membrane (EIHM). At the second infection site no Mep1-mCherry fluorescence could be observed. This suggests breakdown of the EIHM in the second infected cell in which Mep1-mCherry leaks into the whole cell, which loses viability. Laser confocal images were taken at 24 hpi. Scale bars = 10 um.

**Supplemental Figure 12.**
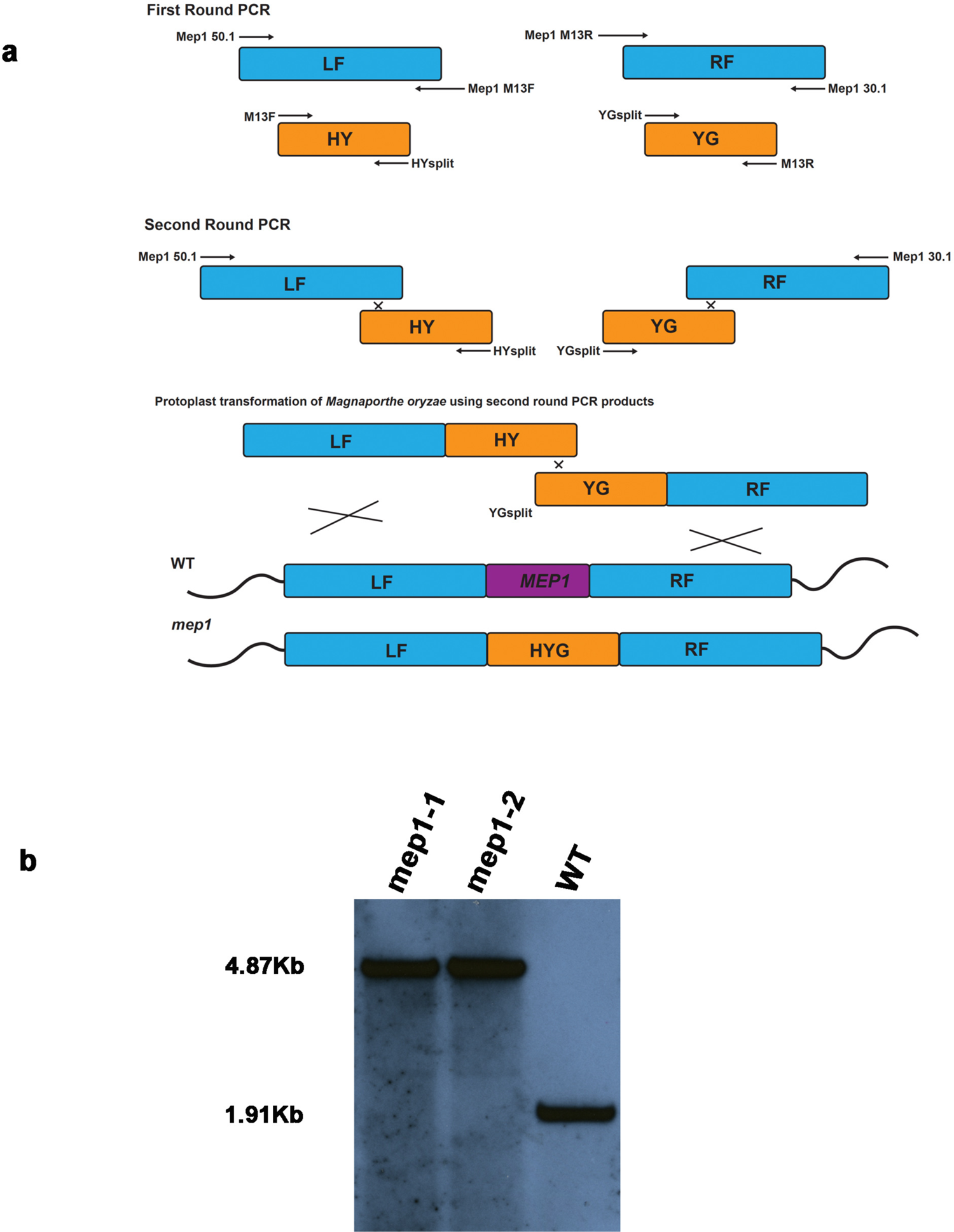
Targeted *MEP1* gene deletion in the rice blast fungus *Magnaporthe oryzae*. **(A)** Schematic representation of the *MEP1* gene deletion strategy using the split-marker strategy, as described in Methods. Two amplicons containing the left and right flanks of *MEP1* and each half of the hygromycin resistance gene cassette are used to transform Guy11. This results in three crossover events to generate a target gene replacement with re-synthesis of the whole hygromycin resistance cassette. **(B)** Southern blot to confirm targeted gene replacement of the *MEP1* gene. The same approach was used to generate gene replacement mutants for each *MEP* gene analysed in this study.

**Supplemental Figure 13.**
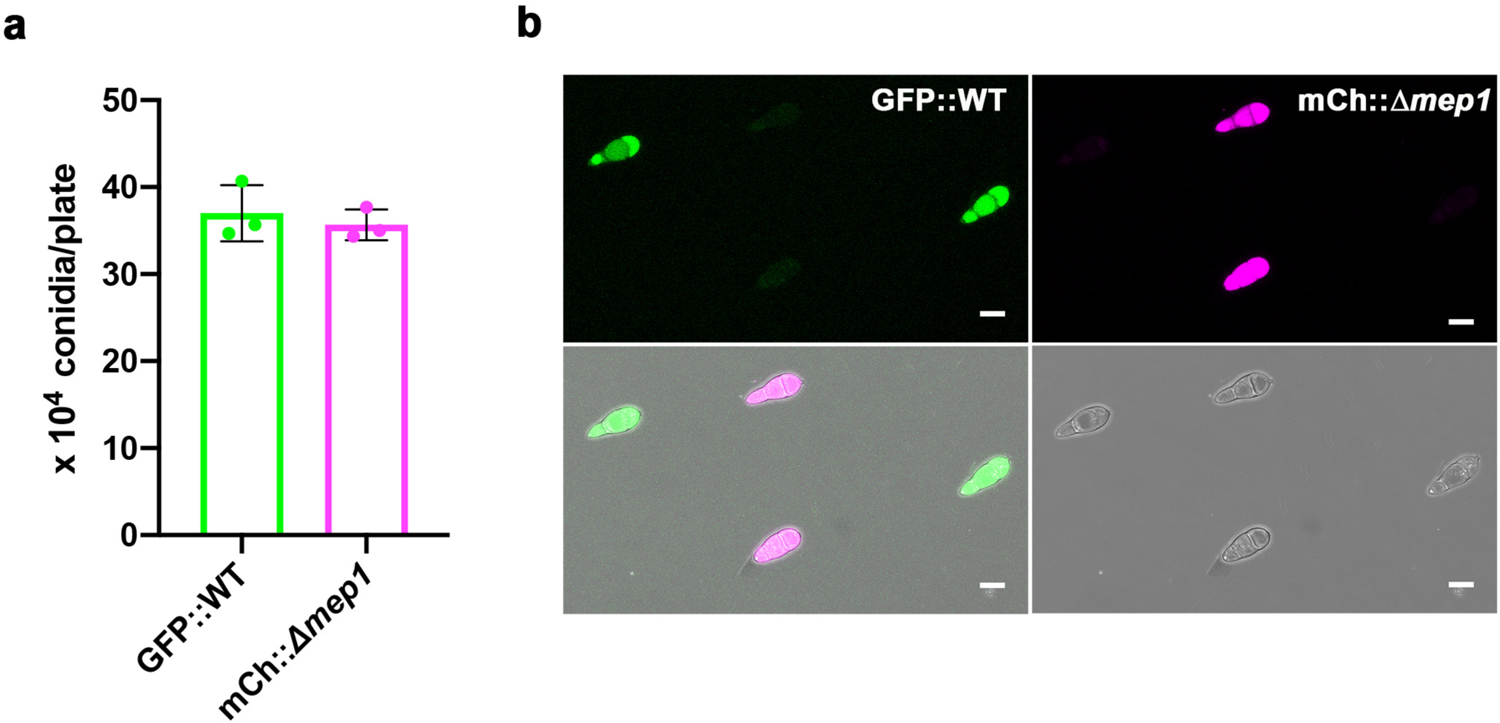
Quantification of conidiogenesis from the *Δmep1* mutant. In order to be able to use a mixed infection fitness assay, it was essential to ensure that each *M. oryzae* strain sporulated with the same frequency **(A)** Conidia were collected from WT::TrpCp:GFP and *Δmep1*::TrpCp:mCherry and counted using a haemocytometer. Conidiation was analysed by quantifying conidia harvested from 10 day-old CM plate cultures of *M. oryzae*. Bar charts show the mean and standard deviation. No significant difference (P > 0.05) in conidiation was observed in three biological replicates of the experiment. **(B)** Laser confocal micrographs showing conidia of WT::TrpCp:GFP and *Δmep1*::TrpCP:mCherry, respectively.

**Supplemental Movie 1.** Real-time observation of the rice blast fungus colonising plant cells. Movie showing the colonisation of rice cells by the *M. oryzae* Guy11 expressing constitutively high level expression vector RP27p:GFP. The starting point of imaging is 38 hpi and images were taken every 30 min for 10 hours before assembling into a single time-lapse movie. Scale bars = 20 μm

**Supplemental Movie 2.** Three dimensional live-cell imaging to show co-localization of Mep1^19-74^-GFP and Mep1-mCherry fluorescence during plant infection.

Movie showing a three-dimensional projection of invasive hyphae of *M. oryzae* expressing Mep1^19-74^-GFP and Mep1-mCherry growing within a living rice cell. Green fluorescence from GFP is uniformly enveloped by the magenta fluorescence from the secreted Mep1-mCherry. Laser confocal mages were taken at 24 hpi. Scale bars = 10 um

**Supplemental Dataset 1.** Summary of read counts in this study.

**Supplemental Dataset 2.** Gene list of members of each WGCNA co-expression module during rice blast disease development.

**Supplemental Dataset 3.** Expression profile of predicted secreted protein-encoding genes differentially expressed during spray-infection of rice cultivar CO39 by *M. oryzae*.

**Supplemental Dataset 4.** Expression profile of predicted secreted protein-encoding genes differentially expressed during spray-infection of rice cultivar Moukoto by *M. oryzae* **Supplemental Dataset 5.** Expression profile of differentially expressed signal peptide containing genes of *M. oryzae* during leaf drop-infection of rice cultivar CO39.

**Supplemental Dataset 6.** List of differentially expressed *MEP* genes that are dependent on the Pmk1 MAP kinase for the differential expression during plant infection.

**Supplemental Dataset 7.** Expression modules and computationally predicted structural clusters of *M. oryzae* secreted proteins.

**Supplemental Dataset 8.** List of *M. oryzae* effector candidates studied by live-cell imaging in this study.

**Supplemental Dataset 9.** Oligonucleotide primers used in this study.

**Supplemental Dataset 10.** Plasmids used in this study.

**Supplemental Dataset 11.** *M. oryzae* strains generated in this study.

